# Allelic coverage analyses reveal a high proportion and uneven chromosomal distribution of SNPs associated with polymorphic duplications in the oyster *Ostrea edulis*

**DOI:** 10.1101/2025.06.24.661118

**Authors:** Lila Colston-Nepali, Sylvie Lapègue, Nicolas Bierne

## Abstract

Oysters are known for their high genetic diversity, potentially high genetic load, and structurally complex genomes. The European flat oyster, *Ostrea edulis*, is no exception. Here, we combined a reference-free analysis of pairs of *k*-mers differing by a single nucleotide (hereafter: het-mers) with a reference-based analysis of allelic coverage using high-coverage short-read sequencing data (70-160X) from five individuals. Up to one-third of SNPs called as heterozygous within an individual exhibited allelic coverage fractions departing from the expected 0.5 ratio. Despite no evidence of recent whole-genome duplication, 40% of het-mers displayed coverage profiles consistent with duplicated genomic segments. We classified AAB het-mers as CN3 and AAAB or AABB het-mers as CN4, corresponding to nucleotide variants located within segments present in three and four copies, respectively, and interpreted as heterozygous and homozygous duplications. These CN3 and CN4 het-mers were typically observed in only a subset of individuals and mostly switched among CN3, CN4, or absence from the het-mer catalog, providing evidence for pervasive paralogous variants associated with rampant polymorphic duplications of genomic segments that vary in copy number among individuals. Reference-based analysis of sequencing depth and allelic coverage fraction similarly showed that SNPs with deviant allelic coverage are frequently associated with duplicated genomic segments segregating within populations, predominantly at low frequencies. Despite their different principles and potential biases, both approaches converged on a minimum estimate of 15% of nucleotide variants being associated with polymorphic duplications. Duplication-associated SNPs occurred in coding and non-coding regions to a similar extent, but their distribution was heterogeneous across chromosomes, with the highest densities on three of the ten chromosomes. These three chromosomes are non-metacentric and particularly prone to chromosome loss in oysters, which are known for their high incidence of somatic aneuploidy. This pattern supports two non-exclusive hypotheses: an architecture-driven model in which chromosome properties prone to structural instability promote both duplications and somatic aneuploidy, and a compensation-driven model in which duplications evolve to buffer dosage imbalance in aneuploid cells. Finally, we discuss how extensive polymorphic duplication may bias population genomic analyses in *O. edulis* and other ostreids.

## INTRODUCTION

Marine bivalves often exhibit very high levels of genetic diversity, making them ideal models for population genetics since the advent of allozyme markers (Ward, Woodwark, and Skibinski 1994; Bazin, Glémin, and Galtier 2006). Early studies revealed several striking observations, including heterozygote deficiencies relative to Hardy-Weinberg equilibrium (Zouros and Foltz 1984), segregation distortion in controlled crosses (Bierne et al. 1998; Launey and Hedgecock 2001), and heterozygosity-fitness correlations (David 1998). These patterns have been interpreted in terms of abundant adaptive variation (Mitton 2000; Koehn 1983), unusual demographic processes such as chaotic genetic patchiness (Eldon et al. 2016; David et al. 1997) and sweepstakes reproductive success (Hedgecock and Pudovkin 2011), high rates of somatic aneuploidy (Thiriot-Quiévreux 1986; Thiriot-Quiévreux et al. 1992; Leitão et al. 2001), and technical issues such as null alleles (Foltz 1986). Very large population sizes may contribute both to high polymorphism (Harrang et al. 2013; Takeuchi 2017) and to a high genetic load (Plough 2016), but not all of these observations have been fully resolved. DNA sequence analyses have confirmed high nucleotide diversity at both silent and amino-acid changing sites in oysters (Sauvage et al. 2007; Harrang et al. 2013; Romiguier et al. 2014). More recently, access to structural variation in genome sequence studies has revived the idea of potentially unique bivalve genetics underpinned by widespread presence-absence variation (Gerdol et al. 2020), copy-number variation (Modak et al. 2021; Qi, Li, and Zhang 2021), and highly divergent haplotypic sequences (Liu et al. 2025), renewing interest in the possibility of unusual genome instability of bivalves. Although this widespread structural variation has been proposed to be adaptive (i.e. maintained by selection) by others, echoing earlier claims for allozyme and SNP variation, we believe that large population sizes and high genetic load, possibly associated with sweepstakes reproduction, may provide a more plausible explanation.

Highly polymorphic species usually have a compact genome (Lynch and Walsh 2007), though this is not always the case, such as in marine pteriomorph bivalves. A high proportion of their genomes are made of repetitive sequences (Šatović and Plohl 2021). This includes Pacific oysters (*Magallana gigas*, Zhang et al. 2012), pearl oysters (*Pinctada fucata*, Takeuchi et al. 2012; Takeuchi et al. 2022), Mediterranean mussels (*Mytilus galloprovincialis*, Gerdol et al. 2020), deep sea and shallow-water mussels (*Bathymodiolus platifrons* and *Modiolus philippinarum*, Sun et al. 2017), scallops (*Patinopecten yessoensis*, Wang et al. 2017), and golden mussels (*Limnoperna fortunei*, Uliano-Silva et al. 2018). Although repetitive sequences include highly repeated elements such as mobile elements, satellite DNA, or tandem repeats (Šatović and Plohl 2021), segmental duplications (SDs) have been found to make up 26% of the Pacific oyster genome, the highest recorded in published genomes to date (Qi et al. 2021). Studies of resequenced genomes have also revealed presence-absence variation (PAV) throughout the Mediterranean mussel genome, with approximately 25% of genes subject to PAV (Gerdol et al. 2020), but also persistent copy-number-variable regions (CNVRs) consisting of over 43% of the Pacific oyster genome (Qi, et al. 2021), and duplication variation contributing to over 16.5% of the eastern oyster (*Crassostrea virginica*) genome (Modak et al. 2021).

Population genomics studies, however, have been predominantly obtained with low coverage genome sequencing in order to analyze many samples within constrained project budgets. These studies focus on SNPs and mostly ignore structural variation, even though some of these SNPs are likely to be found in these parts of the genome. Downstream analyses rely on a number of assumptions about genome assemblies, although next-generation sequencing methods have inherent technical risks of error that may violate these assumptions and lead to misinterpretation of the data. For this reason complex filtering strategies are used to improve the accuracy and reliability of genome-wide SNP datasets (O’Leary et al. 2018; Hemstrom et al. 2024). Very few studies, though, consider the effect of genome complexities on SNP calling in next-generation sequencing data. Dallaire et al. (2023) described the effects of rediploidization and repetitive elements on low-coverage whole genome sequencing in a salmonid fish, which they hypothesize causes collapsed assembled genomic regions. As a result SNPs may be called that do not conform to expected patterns of heterozygosity and allelic coverage fractions, which they refer to as ‘deviant SNPs’. For example, such SNPs deviated from the expected allelic coverage fraction of 1/2, with a hotspot of SNPs demonstrating an allelic coverage fraction of 1/4 (Dallaire et al. 2023). These deviant SNPs have a significant effect on downstream analyses and population genetic inferences, and most filtering approaches will fail to remove them (Dallaire et al. 2023). The presence of these deviant SNPs in non-model species with complex genomes, such as oysters, represents a potential blind spot in population genomic analyses.

The European flat oyster, *Ostrea edulis*, is a marine pteriomorph bivalve from the family Ostreidae, or the true oysters. *O. edulis* has highly variable reproductive success, both in the wild and in experimental conditions, and a low Ne/N ratio (Hedgecock et al. 2007; Lallias et al. 2010). *O. edulis* has been found to have high genetic load causing segregation distortions in pair crosses (Bierne et al. 1998), and up to a third of non-synonymous mutations have been estimated to be mildly deleterious in this species (Harrang et al. 2013). The genome size of *O. edulis* is also larger than other sequenced oyster species (Boutet et al. 2022; Gundappa et al. 2022; Li et al. 2023), and contains comparable repeated elements as *M. gigas*, *C. virginica*, and *Saccostrea glomerata* (Boutet et al. 2022; Li et al. 2023). Synteny analyses indicate that *O. edulis* has not experienced a recent whole genome duplication event since divergence from Crassostreinae (Li et al. 2023), although such an event may have occurred at the origin of all Ostreidae.

The present study was prompted by a highly skewed distribution of allelic coverage fractions in a short-read *O. edulis* dataset initially generated for population genetics. To investigate its origin, we analyzed five oysters sequenced at very high coverage (70-160X) as part of a project initially designed to study somatic mutations. We combined a reference-free analysis of heterozygous *k*-mer pairs (het-mers) using Smudgeplot (Ranallo-Benavidez et al. 2020) with a reference-based analysis of sequencing depth and allelic coverage. The two approaches are complementary: het-mer analysis avoids mapping bias but does not easily localize variants, whereas reference mapping provides genomic positions but can be affected by mapping bias. We first used het-mers to characterize nucleotide variants associated with different copy-number states and to compare these states among individuals. We then used reference-based dosage-inferred genotypes (DI-genotypes) to estimate the frequency and genomic distribution of duplication-associated SNPs. Both approaches converged on a minimum estimate that approximately 15% of nucleotide variants are associated with polymorphic duplications, duplicated genomic segments whose copy number varies among individuals. We finally examined the frequency spectrum, functional annotation, and chromosomal distribution of duplication-associated SNPs, revealing that they are particularly abundant on three of the ten chromosomes, a pattern that had previously gone unnoticed.

## METHODS

### 1. Principle of approaches combining sequencing depth and allelic coverage fraction

Both the het-mer and reference-mapping approaches rely on the joint information provided by sequencing depth and allelic coverage fraction at biallelic nucleotide variants. Under uniform sequencing coverage, total depth is expected to scale with the number of genomic copies contributing reads to a locus, whereas the relative coverage of the two nucleotide variant alleles reflects their dosage among these copies. Thus, a standard diploid heterozygous site (AB) is expected at 2X the average haploid depth (CN2) with an allelic coverage fraction of 1/2; an AAB configuration is expected at 3X depth (CN3) with a minor-allele fraction of 1/3; and AAAB and AABB configurations are expected at 4X depth (CN4) with minor-allele fractions of 1/4 and 1/2, respectively (Fig. 1). Homozygous configurations such as AA, AAA, and AAAA (or BB, BBB, BBBB) have the corresponding CN2, CN3, and CN4 depths but an allelic coverage fraction of zero. Het-mer analysis is reference-free and therefore avoids mapping bias but does not easily provide the genomic position of the variant. Conversely, reference mapping localizes variants but relies on reads originating from different copies mapping to the same location in the reference genome. This is expected when the reference genome carries the 1-copy allele and additional copies in CN3 or CN4 individuals are sufficiently similar to map back to that locus. If the reference instead contains the duplicated state, reads may map to multiple paralogous positions and be removed by standard mapping-quality filters. Importantly, our reference-based approach detects excess copy number relative to the reference but does not identify the physical location of the additional copy, which may be tandem, elsewhere on the same chromosome, or on another chromosome.

**Figure 1.**
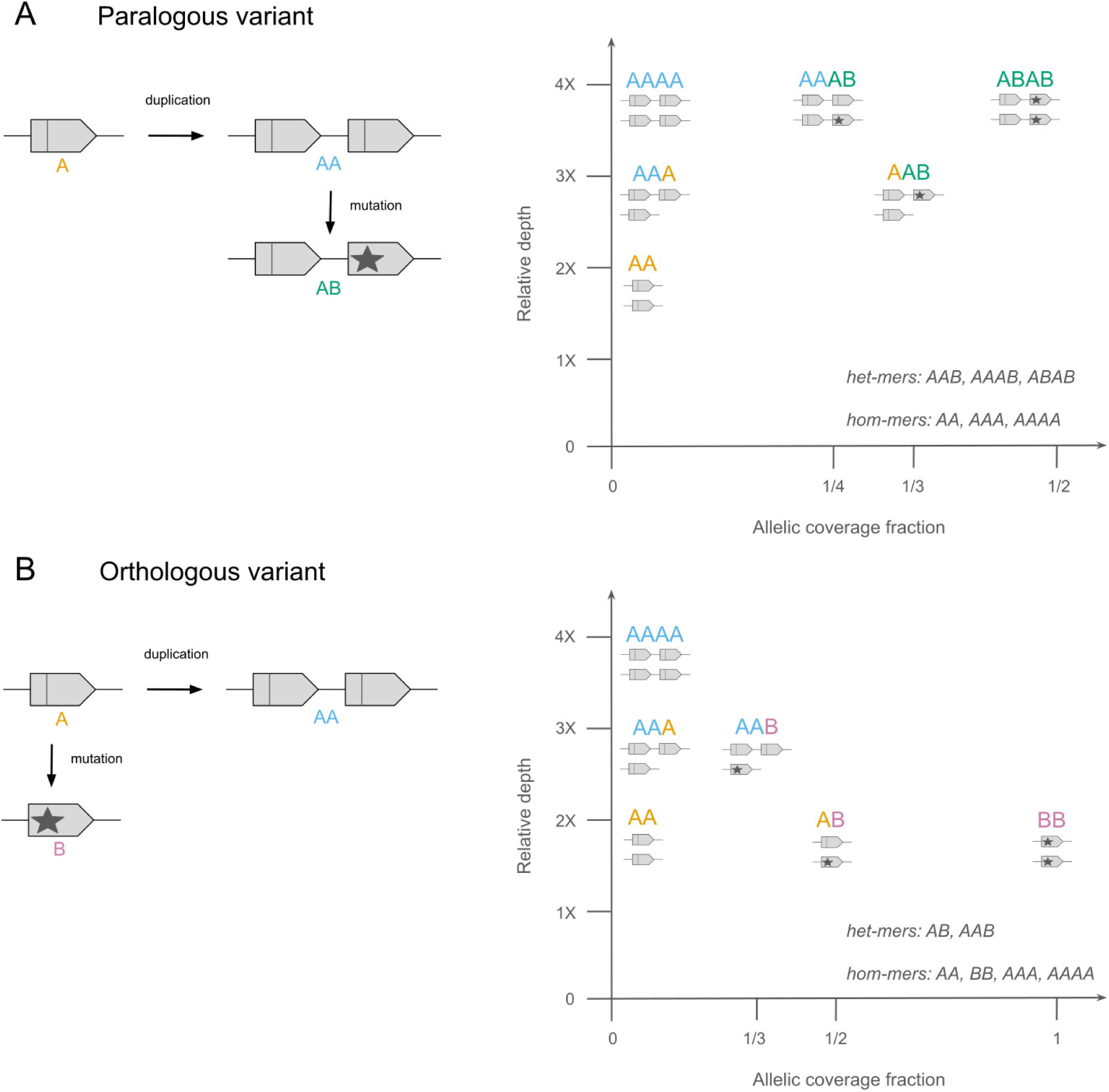
Principle of copy-number inference from the joint distribution of sequencing depth and allelic coverage fraction. Two simple mutational paths illustrate how nucleotide variants located within polymorphic duplicated segments generate distinct combinations of relative depth and allelic coverage fraction.

To interpret these configurations in the context of polymorphic duplications, we distinguish between nucleotide variants at a SNP and copy-number alleles of the genomic segment containing that SNP. We consider a 1-copy allele as a haplome carrying one copy of the focal segment and a 2-copy allele as a haplome carrying two copies, irrespective of the physical location of the additional copy within that haplome. Here, we use haplome to refer to one of the two parental haploid genomic complements of a diploid individual, without implying that the additional copy is located on the same chromosome as the original copy. A duplication initially produces a 2-copy allele whose two copies are identical and therefore does not, by itself, generate a nucleotide variant detectable as a het-mer or SNP. A subsequent nucleotide mutation can generate such a variant through two simple mutational paths that differ in whether the nucleotide variation occurs between paralogous or orthologous copies (Fig. 1). In the first path, which we refer to as the paralogous-variant path (Fig. 1A), duplication occurs first and a nucleotide mutation subsequently arises in one of the two paralogous copies of the 2-copy allele. Starting from an ancestral A sequence, duplication generates a 2-copy AA haplotype, and mutation in one copy subsequently generates a 2-copy AB haplotype. If not lost by genetic drift, three segment haplotypes can therefore segregate in the population: a 1-copy A haplotype, an unmutated 2-copy AA haplotype, and a mutated 2-copy AB haplotype. Their diploid combinations generate configurations including AA, AAA, AAB, AAAA, AAAB, and AABB genotypes. Critically, the A/B nucleotide difference exists only between the two paralogous copies of the duplicated allele: there is no 1-copy B haplotype and therefore no standard CN2 heterozygous AB genotype. Consequently, when reads from the two paralogous copies are mapped to the same reference position, their fixed or segregating difference can be called as a heterozygous SNP even though it does not represent allelic variation between orthologous copies. This is the situation with the greatest potential to affect population genetics analyses, because a paralogous sequence variant may be treated as a conventional heterozygous SNP, and its frequency will primarily reflect the frequency of the duplicated segment rather than that of an orthologous nucleotide allele. In the second path, the orthologous-variant path (Fig. 1B), the nucleotide mutation occurs on a 1-copy haplotype rather than between the two copies of the duplicated segment. Starting from a 1-copy A allele, the population can therefore contain the standard 1-copy A and B alleles, together with the duplicated 2-copy AA haplotype. The A/B nucleotide polymorphism is thus a conventional orthologous polymorphism, and combinations of the 1-copy A and B alleles generate the standard CN2 genotypes AA, AB, and BB. When the 2-copy AA haplotype is present, it contributes two A copies and generates CN3 or CN4 configurations such as AAA, AAB, and AAAA. In this case, the additional paralogous copy modifies total depth and, in heterozygous configurations such as AAB, shifts the expected allelic coverage fraction from 1/2 to 1/3. However, unlike in the paralogous-variant path, the A/B SNP itself represents genuine orthologous variation and can also occur as a conventional AB genotype at CN2. The main consequence of the duplication for standard population genetics analyses is therefore distortion of allelic coverage and potentially genotype likelihoods or genotype calling, rather than the creation of an apparent SNP that does not exist as an orthologous polymorphism.

These two simple paths illustrate why nucleotide variants can act as markers of the copy-number state of the genomic segments in which they occur, while also showing that duplication-associated SNPs need not have the same origin. The distinction cannot always be established from sequencing depth and allelic coverage alone, and the configurations in Fig. 1 should therefore be regarded as parsimonious models rather than unique evolutionary histories. Other scenarios are possible, including loss of one copy from an ancestrally duplicated segment, recombination between duplicated copies, or secondary contact between lineages differing in copy number and sequence divergence. Likewise, sequencing depth provides the total number of copies contributing reads but does not determine how these copies are partitioned between the two haplomes. For subsequent analyses, we therefore assumed that CN2, CN3, and CN4 correspond to 1+1, 1+2, and 2+2 copies across the two haplomes, respectively, and thus to zero, one, and two 2-copy segment alleles.

Alternative configurations, such as 0+3 for CN3 or 1+3 and 0+4 for CN4, cannot be excluded from sequencing depth and allelic coverage alone but were here assumed to be sufficiently rare to neglect them.

### 2. Sampling and sequencing

Five *O. edulis* samples with high-coverage short-read sequencing were used: four newly sequenced samples from Galicia, Spain (approximately 70-160X), and one from Brittany, France, using publicly available reads (SRA accession SRR1723031; ∼70X; named ROSC in this study; abductor muscle tissue). Three Galician samples (2C4M, 6C9M, and 8C17M) came from a culture raft in Cambados, Ría de Arousa, and one sample (4C6M) came from a natural bed in Ría de Ferrol. DNA was extracted from mantle tissue using the Macherey-Nagel NucleoSpin Tissue kit. TruSeq DNA PCR-Free libraries were prepared and sequenced at the MGX platform on an Illumina NovaSeq 6000 S4 flow cell.

### 3. Reference-free het-mer analysis

We first used a reference-free *k*-mer based analysis. *k*-mer approaches allow users to evaluate large, and potentially complex, genomic datasets computationally efficiently and without the need for a reference genome (Jenike et al. 2025). The program Smudgeplot (Ranallo-Benavidez et al. 2020) uses the co-distribution of total coverage and the minor *k*-mer coverage of pairs of *k*-mers differing by a single nucleotide to determine how many copies of each *k*-mer exist across the genome. Therefore, Smudgeplot allows the delineation of distinct dosage clusters that would be otherwise difficult to determine using coverage alone (even when coverage is high, as is the case here). Smudgeplot analyses are completed on individual genomes.

First, *k*-mer frequency histograms were computed for each individual using the program FastK (https://github.com/thegenemyers/FASTK) on the raw reads, using a *k*-mer count of 21. Then, the program GenomeScope 2.0 (Ranallo-Benavidez et al. 2020) was used to analyze the *k*-mer coverage for each individual, and fit to a model of 2 x *p* negative binomial distributions (*p* = ploidy) representing the unique and multi-copy heterozygous and homozygous *k*-mers. A ploidy of *p* = 2, or diploid, was used. GenomeScope 2.0 was used to determine input parameters for error margins and coverage ranges in the execution of Smudgeplot. Next, the program Smudgeplot v. 0.5.4 (Ranallo-Benavidez et al. 2020) was used to find *k*-mer pairs that differ by exactly one nucleotide (heterozygous *k*-mers, or het-mers), representing either true heterozygous alleles of orthologous segments, or nucleotide variants associated with complex genome structures combining orthologous and paralogous segments that vary in copy number, using the function -*hetmers*. Total coverage of the het-mers was plotted against the minor *k*-mer coverage to reveal hotspots that represent the haplotype structures present, and their frequency, in the genome using the function -*all*. Custom parameters for each individual for these functions were determined using the GenomeScope 2.0 results. Unlike GenomeScope 2.0, ploidy is not needed a priori to run Smudgeplot.

Finally, to determine whether the same nucleotide variants represented by Smudgeplot het-mers were shared among individuals and whether their inferred copy-number states varied among individuals, we used the Smudgeplot -*extract* function to retrieve the sequences of the *k*-mer pairs assigned to each smudge category. Custom Python scripts were then used to match identical pairs of *k*-mers across individuals and generate a comparison table reporting the Smudgeplot category assigned to each marker in each individual. We considered AB het-mers to represent the standard diploid CN2 state, AAB het-mers to represent CN3, and AAAB or AABB het-mers to represent CN4. These categories describe the dosage of the two nucleotide variants among the inferred number of copies. Importantly, Smudgeplot identifies only pairs of *k*-mers differing by a single nucleotide and therefore detects a marker only when both nucleotide variants are present in an individual. Consequently, a marker identified as a het-mer in one individual may not be recovered in another individual if the corresponding nucleotide site is invariant, irrespective of its copy-number state. This includes standard homozygous CN2 genotypes (AA or BB), as well as invariant CN3 or CN4 configurations (AAA, BBB, AAAA, or BBBB). When a marker identified in at least one individual was not recovered as a het-mer in another individual, it was assigned to the category ABS (absent), indicating absence from the individual’s het-mer catalog rather than an inferred copy-number state. For each marker, the combination of states observed across the five individuals (AB, AAB, AAAB/AABB, or ABS) was recorded. To summarize the patterns independently of individual identity, the combinations of states observed across the five individuals were sorted alphabetically before counting their frequencies, thereby treating combinations differing only in the identity of the individual carrying a given state as equivalent.

In order to investigate and compare genome-wide duplication events in other ostreids, Smudgeplot analyses were also performed on publicly available short-read datasets of three oyster species: the lamellated oyster *O. denselamellosa* (SRA accession: SRR19238449), the Olympia oyster *O. lurida* (SRA accessions: SRR3306651-SRR3306656), and the Pacific oyster *M. gigas* (SRA accession: SRR7474375). The SRA toolkit (https://github.com/ncbi/sra-tools) was used to download and extract FASTQ files. For *O. lurida*, six sequencing runs of the same individual were pooled.

### 4. Reference-based analyses

Reference-free het-mer analysis identifies nucleotide variants in segments that vary in copy number, but these markers cannot be directly equated with SNPs called after reference mapping in population genomic studies and are not readily localized in the genome. We therefore used a complementary reference-mapping approach to characterize the genomic distribution of duplication-associated SNPs.

#### 4.1 Alignment to reference genome and SNP calling

Sequence data was trimmed using the program fastp (Chen et al. 2018) to filter for quality and remove both polyG tails and Illumina adapters. Samples were aligned to an *O. edulis* reference genome (Li et al. 2023, GenBank assembly: GCA_032173915.1) using the program BWA-MEM (Li and Durbin 2010). Mapped reads were then filtered using the program Sambamba (Tarasov et al. 2015) to remove reads that did not pair properly, and secondary alignments. SNPs were called using bcftools mpileup and bcftools call (Danecek et al. 2021), with maximum coverage increased to 2000, resulting in 20,868,611 nucleotide variants.

#### 4.2 Dosage based inference of genotypes

Output VCF files were filtered to retain only biallelic SNPs and to remove indels and sites with missing data using VCFtools (Danecek et al. 2011) and BCFtools (Danecek et al. 2021). SNPs mapping to unplaced scaffolds were removed in R (R Core Team 2023), leaving 15,106,345 nucleotide variants for downstream analyses.

Inspired by Smudgeplot, we used reference-aligned reads and called SNPs to infer dosage-based genotypes (hereafter DI-genotypes) from the joint distribution of sequencing depth and allelic coverage fraction. A custom Python script was used to extract the read counts supporting the reference (A) and alternative (B) alleles at each SNP, calculate total allelic depth (A+B), and estimate the allelic coverage fraction for each of the five individuals. Allelic coverage fraction was defined as the proportion of reads supporting the alternative allele, B/(A+B). The resulting CSV files were analyzed in R (R Core Team 2023).

First, distributions of allelic coverage fractions at heterozygous SNPs were examined using histograms for each individual. Histograms of total allelic depth were then used to estimate individual average sequencing depth and determine the expected depth for the standard diploid (CN2) state from the mode of the distribution. These estimates were consistent with the initial sequencing coverage estimates. Based on the estimated CN2 depth, the expected depths corresponding to CN1, CN3, and CN4 states were calculated for each individual. Higher copy-number states (CN5 and above) were excluded from subsequent analyses because they represented only a small proportion of SNPs.

Density plots were generated using ggplot2 (Wickham 2016) to visualize the joint distribution of total allelic depth and allelic coverage fraction. Based on their expected positions in this two-dimensional distribution, DI-genotype clusters were defined and stringently circumscribed for 12 possible genotypes at biallelic sites: AA, AB, BB, AAA, AAB, ABB, BBB, AAAA, AAAB, AABB, ABBB, and BBBB. Unlike the het-mer approach, reference-based analysis also recovers genotypes in which only one nucleotide variant is present (AA, AAA, AAAA and BB, BBB, BBBB), which are absent from the het-mer catalog. This allows a more complete characterization of genotype frequencies and, under our 1-copy/2-copy segment allele interpretation, of the frequency of the corresponding segment alleles. For example, a standard heterozygous diploid genotype (AB) is expected to combine CN2 depth with an allelic coverage fraction of 0.5, whereas AAB and ABB genotypes combine CN3 depth with expected allelic coverage fractions of 1/3 and 2/3, respectively. Similarly, AAAB, AABB, and ABBB combine CN4 depth with expected allelic coverage fractions of 1/4, 1/2, and 3/4, respectively. To minimize ambiguous assignments between neighboring clusters, DI-genotypes were assigned only to SNPs falling within regions of the joint distribution for which genotype inference was considered reliable. This high-confidence subset was retained for subsequent analyses of variation among individuals, chromosomes, and functional categories. Finally, DI-genotypes were grouped according to inferred copy number: AA, AB, and BB were classified as CN2; AAA, AAB, ABB, and BBB as CN3; and AAAA, AAAB, AABB, ABBB, and BBBB as CN4.

#### 4.3 Location and length distribution of runs of CN3 and CN4 SNPs

To investigate the genomic distribution of SNPs associated with duplicated segments, the high-confidence DI-genotypes described above were first positioned along each of the ten chromosomes for each of the five individuals. SNPs were classified as CN2, CN3, or CN4 according to their DI-genotype, and their genomic positions were used to visualize the distribution of the three copy-number states along chromosomes. This chromosome-wide representation allowed us to assess variation among individuals and heterogeneity in the occurrence of CN3 and CN4 SNPs along and among chromosomes.

Multiple neighboring SNPs assigned to CN3 or CN4 are expected when they occur within the same duplicated genomic segment, so we then identified runs of SNPs showing consistent CN3 or CN4 dosages. Runs were detected independently for CN3 and CN4, for each individual and chromosome. A run was required to start with a SNP of the target copy-number state, to contain at least five SNPs, and to be at least 150bp long. To account for occasional discordant DI-genotype assignments within an otherwise consistent region, up to 10% of SNPs within a run were allowed to have a copy-number state different from the target state (note that this means no inconsistent SNPs for runs smaller than 10 SNPs). A run was interrupted when this proportion was exceeded or when two consecutive SNPs were separated by more than 100 kb. Non-target SNPs occurring at the end of a candidate run were excluded so that the boundaries of each run corresponded to target CN3 or CN4 SNPs. The physical length of each run was calculated as the distance in base pairs between its first and last target SNP.

#### 4.4 Segmental duplication allele frequency spectrum

To determine whether the same SNPs were assigned to the same DI-genotypes across individuals, SNP lists were generated for each of the 12 DI-genotype categories in each individual. These lists were then combined into a comparison table reporting the DI-genotype assigned to each SNP in each of the five individuals. We subsequently used these DI-genotypes to infer the frequency of the duplicated genomic segments in which the SNPs are located. Importantly, we distinguish here between the nucleotide alleles (A and B) at a SNP and the copy-number alleles of the genomic segment containing that SNP. For the latter, we considered two segment alleles: a 1-copy allele, corresponding to a haplome carrying a single copy of the segment, and a 2-copy allele, corresponding to a haplome carrying a duplicated copy of the segment. To infer segment copy-number genotypes, we assumed that CN2 corresponds to two 1-copy segment alleles (1-copy/1-copy), CN3 to a heterozygous duplication carrying one 1-copy and one 2-copy segment allele (1-copy/2-copy), and CN4 to a homozygous duplication carrying two 2-copy segment alleles (2-copy/2-copy). Under this model, CN2, CN3, and CN4 individuals therefore carry zero, one, and two 2-copy segment alleles, respectively. Each DI-genotype was then recoded according to the inferred number of 2-copy segment alleles as 0 for CN2, 1 for CN3, and 2 for CN4. These values were summed across the five individuals for each SNP, yielding a 2-copy segment allele count ranging from 0 to 10. Thus, although each SNP was used as a marker to infer copy number, the resulting count refers to the frequency of the 2-copy allele of the genomic segment containing the SNP, rather than to the frequency of either nucleotide allele at the SNP itself. The distribution of these counts across SNPs was used to construct a 2-copy segment allele frequency spectrum. A count of 0 indicates that all five individuals were inferred to carry two 1-copy segment alleles at that SNP position (CN2), whereas a count of 10 indicates that all five individuals were inferred to carry two 2-copy segment alleles (CN4). A count of 5 can result either from all five individuals being heterozygotes for the segment duplication or more likely from a combination of CN2, CN3, and CN4 genotypes. All other intermediate counts necessarily indicate variation in inferred segment copy number among individuals.

#### 4.5 Functional annotation of SNPs and functional enrichment analysis

SnpEff (Cingolani et al. 2012) was used to functionally annotate SNPs. A custom SnpEff database for *O. edulis* was built using the genome annotation, coding sequence (CDS), and protein sequence files corresponding to the reference genome assembly used for read mapping (Li et al. 2023). SnpEff annotation categories were simplified into four main classes: intergenic, intronic, synonymous, and non-synonymous variants (see Supplementary Table 1 for details). The proportions of SNPs assigned to CN2, CN3, and CN4 DI-genotypes were then calculated within each annotation category to assess whether duplication-associated SNPs were differentially represented among coding and non-coding regions. To investigate whether the heterogeneous chromosomal distribution of CN3 and CN4 SNPs could be related to differences in gene content among chromosomes, functional enrichment analyses were performed separately for each chromosome using the KEGG Orthology-Based Annotation System (KOBAS; Bu et al. 2021). *Magallana gigas*, the most closely related species available in KOBAS, was used as the reference species. Enrichment was tested using Fisher’s exact test, and P-values were corrected for multiple testing using the Benjamini-Hochberg false discovery rate procedure (Benjamini and Hochberg 1995).

## RESULTS AND DISCUSSION

We first describe the skewed distribution of allelic coverage fractions that prompted this study, then use complementary reference-free het-mer and reference-based approaches to identify SNPs associated with polymorphic duplications. We subsequently examine their frequency and genomic distribution.

### 1. Allelic coverage fractions are highly skewed in O. edulis

We first examined the allelic coverage fraction of SNPs called as heterozygotes in individuals and found that a large proportion (approximately 30%) had a coverage fraction between 0.2 and 0.4 (Fig. 2 and Supplementary Figure 1). Routine inspection of allelic coverage is useful for quality control of next-generation sequencing (NGS) data (Ament-Velásquez et al. 2016; Jaron et al. 2022; Bierne 2022), yet such distributions are still rarely reported. The magnitude of the deviation observed here is unusual. Presence-absence variation, which produces hemizygosity, cannot by itself explain this pattern because it should generate homozygous calls in runs of homozygosity rather than a shift in allelic coverage in heterozygote calls. Plausible explanations instead include paralogy caused by fixed duplications collapsed in the reference or by polymorphic duplications, mapping bias, and somatic aneuploidy. Distinguishing these possibilities requires combining allelic coverage with relative sequencing depth: paralogy is expected to increase depth above the diploid 2X state and make allelic coverage fractions very close to 1/4, 1/3 or 1/2, whereas mapping bias or chromosome loss should reduce depth and make allelic coverage fractions below 1/2 but not necessarily close to 1/4 or 1/3.

### 2. Het-mer analysis reveals a high proportion of CN3 and CN4 het-mers in O. edulis and other oyster genomes

**Figure 2.**
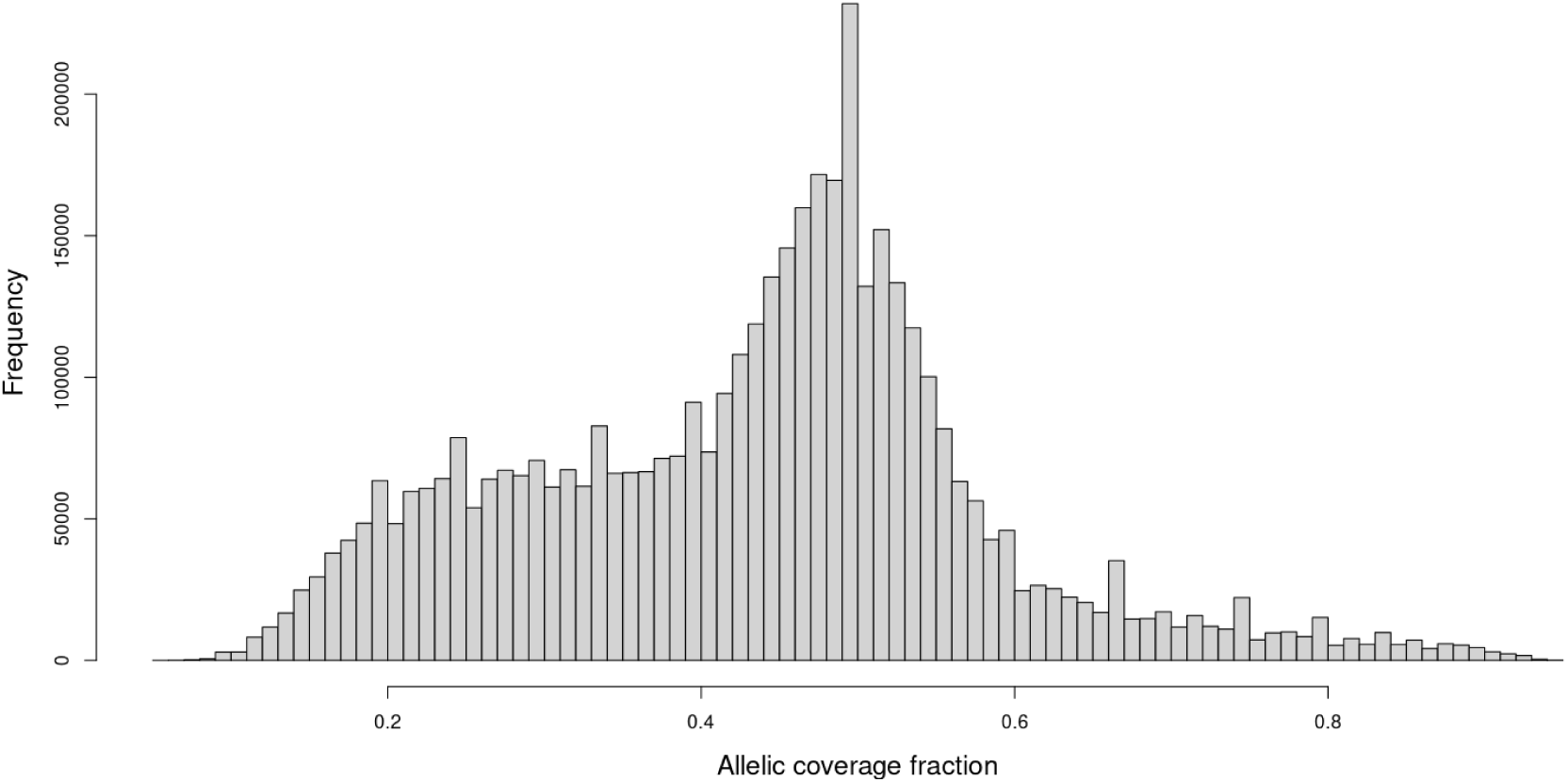
Distribution of the allelic coverage fraction for heterozygote calls in individual 6C9M.

The het-mer coverage distributions obtained with Smudgeplot show that a substantial proportion of het-mers (> 50%) are classified outside the standard CN2 coverage expectations in the genomes of all five individuals (Fig. 3 and Supplementary Figures 2-5), with a high proportion of CN3 and CN4 het-mers. A small proportion of het-mers were assigned to CN5 and higher categories, but these were not considered further because of their very low incidence. Smudgeplot analyses are consistent with widespread duplicated genomic segments, with 14–16% of het-mers assigned to the CN3 category (AAB) and 21–26% assigned to the CN4 categories (AAAB or AABB). The AAB category was particularly informative because it is not expected in a diploid genome except when one haplome carries a single-copy segment allele and the other carries a duplicated segment, as expected for a heterozygous segmental duplication.

**Figure 3.**
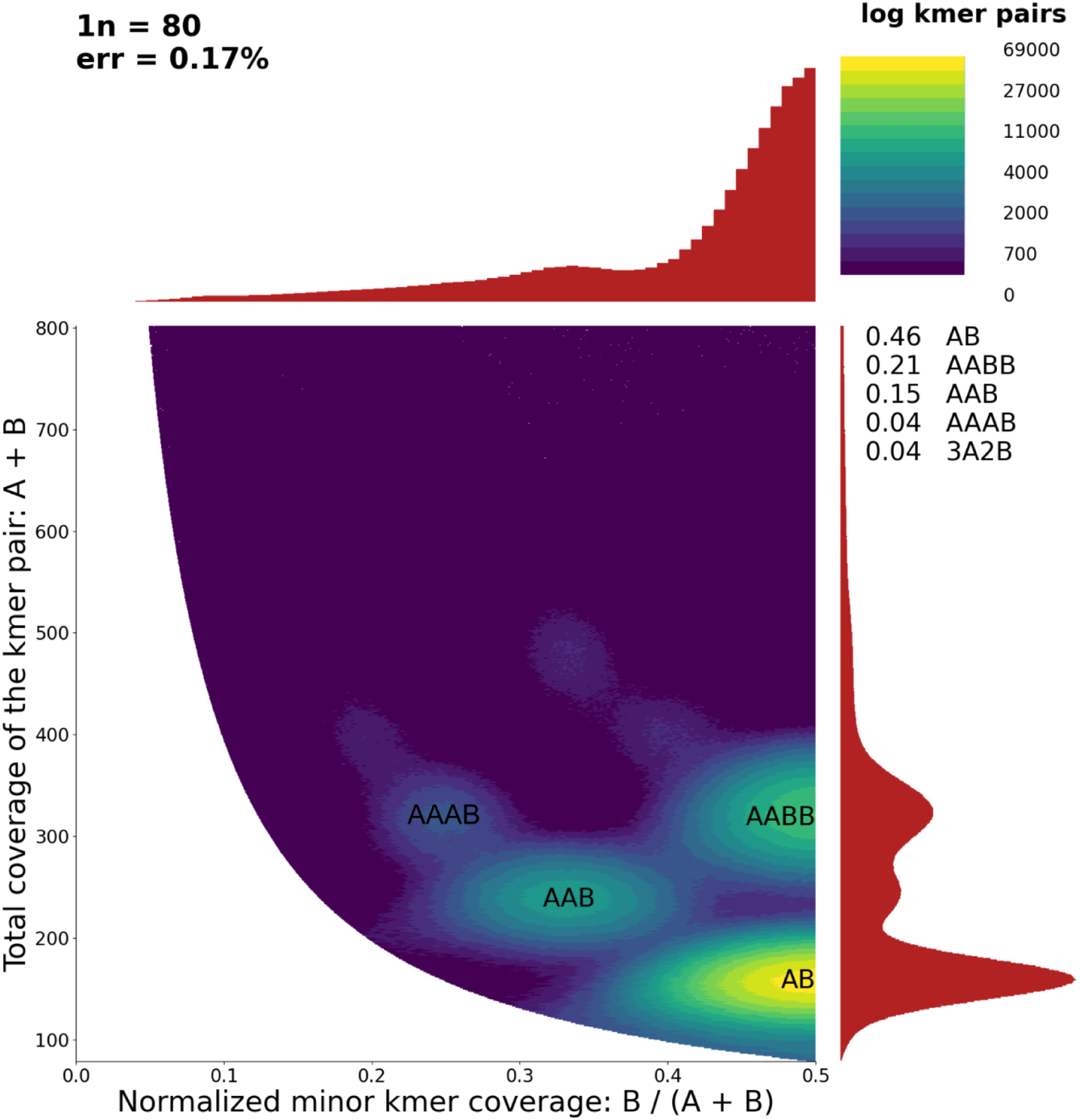
Smudgeplot analysis for sample 6C9M, using an error threshold of 40 and a coverage range of 50-110.

When Smudgeplot markers were compared among individuals, the largest category (66.3% of all markers) comprised markers displaying only the standard CN2 state (AB) and ABS (absent from the het-mer catalog) across individuals (Supplementary Table 2). Within this category, the most common pattern consisted of one AB individual and four ABS individuals (40.2%). These markers most likely correspond to standard orthologous nucleotide variants segregating in diploid genomic segments, with the ABS state resulting from homozygous genotypes (AA or BB) that are not detected by Smudgeplot. Only 2.4% of markers were classified as CN4 (AAAB or AABB) in all individuals, consistent with nucleotide variants associated with fixed duplicated segments. An additional 12.3% of markers displayed a CN4 state in at least one individual but ABS in all others. Because no individual exhibited the heterozygous CN3 (AAB) state, these patterns cannot be unambiguously attributed to polymorphic duplications. They are equally compatible with fixed duplicated segments for which some individuals carry homozygous CN4 genotypes (AAAA or BBBB) that are not detected by Smudgeplot. Consequently, 14.7% represents the maximum proportion of markers that could be associated with fixed duplicated segments.

Only 2.1% of markers combined a CN2 state (AB) with a CN3 (AAB) state, consistent with orthologous nucleotide variants within polymorphic duplicated segments (Fig. 1B). In contrast, 15.9% of markers exhibited at least one CN3 state (AAB genotype) but were never observed in combination with the AB genotype, consistent with the prediction for paralogous nucleotide variants associated with polymorphic duplications (Fig. 1A). These markers therefore provide evidence for a high incidence of paralogous variants associated with polymorphic duplicated segments in *O. edulis*. Within this category, the most frequent pattern consisted of one AAB individual and four ABS individuals (9.1% overall, representing 57% of the category), indicating that many polymorphic duplicated segments are detected as heterozygous duplications in only one individual of our sample. However, a similarly high proportion of singletons was observed among standard CN2 markers: 1 AB + 4 ABS genotypes represented 40.2% overall, or 60% of the 66.3% CN2 category. Thus, the predominance of singleton AAB genotypes largely mirrors the frequency distribution of standard nucleotide polymorphism in our sample. Only 1% of het-mer markers showed more complex configurations combining AB, AAB, and AAAB/AABB genotypes, which cannot be explained by the simple mutational pathways illustrated in Figure 1.

Given that Smudgeplot has rarely been applied to oyster genomes, we reanalysed several public NGS datasets from ostreids (*Ostrea denselamellosa*, *O. lurida*, and *Magallana gigas*) for comparison with *O. edulis*. CN3 (AAB) and CN4 (AAAB and AABB) het-mers were detected in every genome analyzed, but *O. edulis* exhibited the highest proportion of CN3 and CN4 het-mers among the species examined here (Supplementary Figures 6–8). Although *M. gigas* has previously been reported to possess abundant copy-number variable regions and segmental duplications (Qi et al. 2021, 2023), it showed the lowest proportion of CN3 and CN4 het-mers among the ostreids analysed in this study. These observations suggest that Smudgeplot deserves wider application as a routine tool for investigating polymorphic duplicated genomic segments.

Reference-free het-mer analysis therefore provides a first indication of a high incidence of nucleotide variants associated with polymorphic duplications in *O. edulis*. However, a complementary reference-based analysis of SNP allelic dosage was required to assess the consequences for population genomic analyses and to characterize the genomic distribution of CN3- and CN4-associated SNPs among functional categories and along and across chromosomes.

### 3. Reference mapping analysis confirms a high incidence of SNPs associated with polymorphic duplications in O. edulis

In order to better delineate CN3 and CN4 associated SNPs, we adopted a similar approach to Smudgeplot by investigating the co-distribution of relative depth and allelic coverage fraction of heterozygote calls after mapping on a reference genome. SNPs adhering to the expectations for heterozygous diploid sites, AB sites (i.e. CN2 relative depth and 0.5 allelic coverage fraction), had the strongest density (Fig. 4; Supplementary Figures 9-12) as expected.

**Figure 4.**
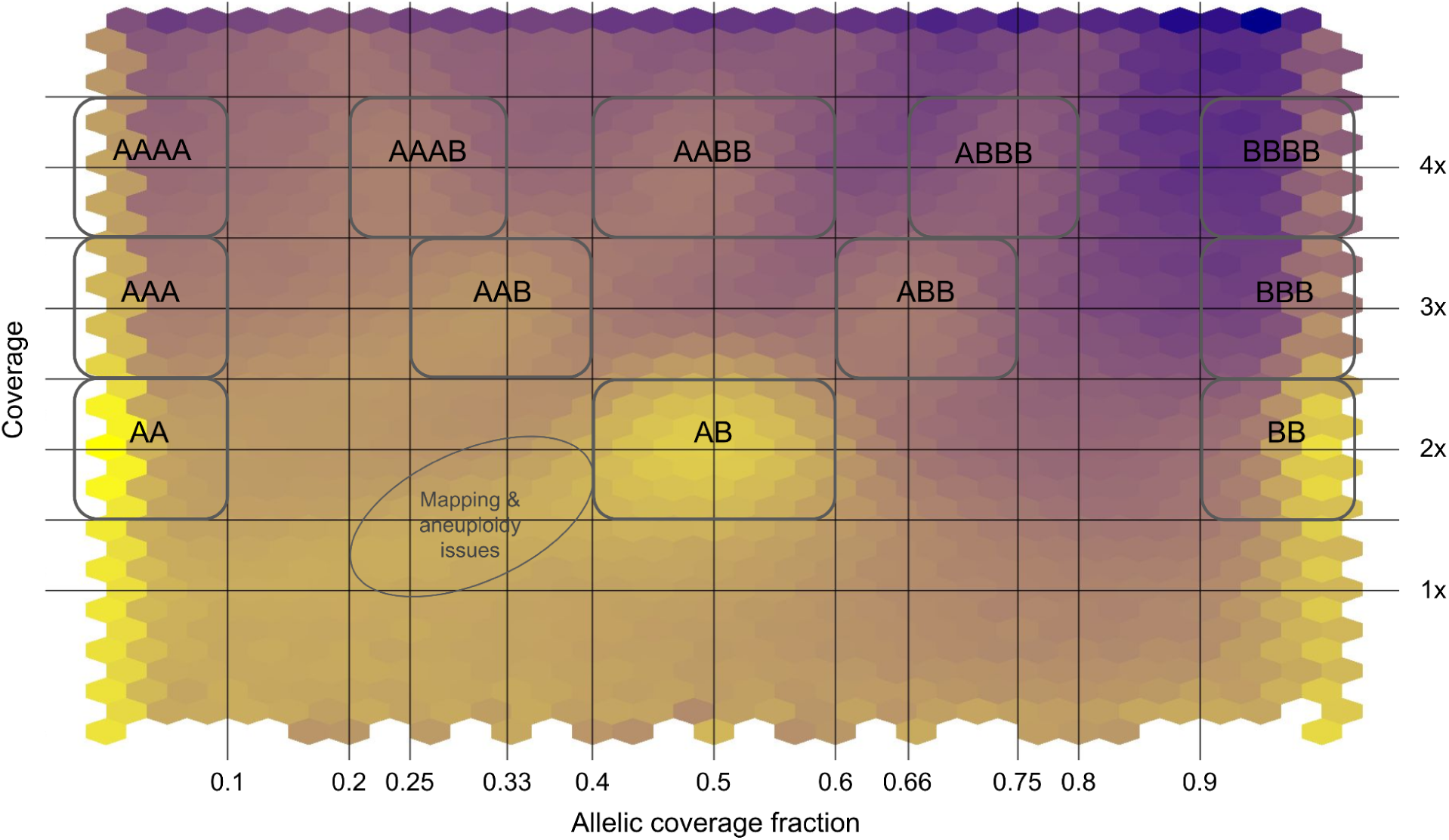
Density plot for sample 6C9M showing the joint distribution of allelic coverage fraction (x-axis) and relative sequencing depth (y-axis). Relative depth corresponds to SNP coverage normalized by the estimated haploid genome coverage, with the expected CN2, CN3 and CN4 depths indicated as 2X, 3X and 4X, respectively. Higher SNP densities are shown in brighter yellow. Boxes delineate the regions used to assign dosage-inferred genotypes (DI-genotypes) based on their expected combinations of relative depth and allelic coverage fraction. For example, the standard diploid heterozygous genotype AB is expected at 2X depth and an allelic coverage fraction of 0.5, whereas AAB and AAAB are expected at 3X and 1/3 and at 4X and 1/4, respectively. The oval indicates a region of increased SNP density that does not correspond to an expected DI-genotype of duplication-associated SNPs and may result from mapping bias or somatic aneuploidy.

Corroborating the Smudgeplot results, strong density was also observed in DI-genotype clusters corresponding to CN3 and CN4 states (Fig. 4; Supplementary Figures 9-12). We additionally observed a region with relative depth below 2X and allelic coverage fraction below 0.5, as expected when one allele maps less efficiently or when a fraction of cells is monosomic, consistent with the high incidence of somatic aneuploidy reported in oysters (Leitão et al. 2001). We did not analyze this lower-depth signal further here and instead focus on the well-defined CN3 and CN4 clusters associated with increased copy number.

The 2-copy segment allele frequency spectrum inferred from DI-genotypes showed that 85% of SNPs were standard CN2 SNPs (2-copy segment allele count = 0; Fig. 5), whereas 15% were associated with duplicated segments with a 2-copy segment allele count greater than zero. Most of these duplication-associated SNPs had counts of 1 or 2, corresponding respectively to one CN3 individual, or to either two CN3 individuals or one CN4 individual among the five sampled oysters. Fixed duplications (count = 10) were rare (0.2%), as expected because the reference-based approach detects polymorphic duplications when the reference carries the 1-copy segment allele. Fixed duplications should generally be absent unless duplicated copies were collapsed during reference assembly. Thus, the reference-based DI-genotype analysis independently supports the het-mer result, indicating that at least 15% of SNPs are associated with polymorphic duplications. Most of these duplications segregate at low frequency, as typically observed for putatively neutral genetic variants.

**Figure 5.**
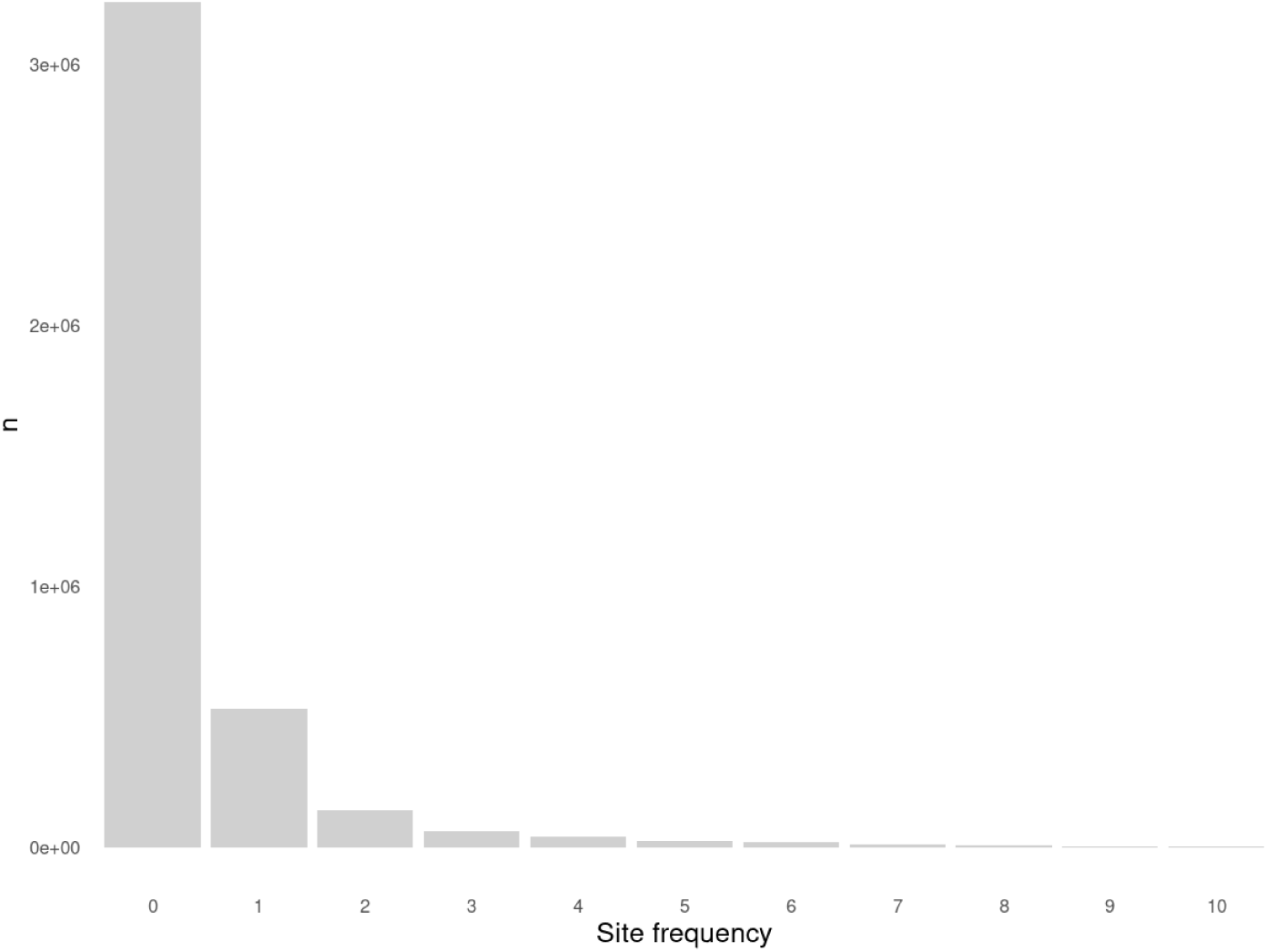
2-copy segment allele frequency spectrum obtained from DI-genotypes across five *O. edulis* individuals.

### 4. Duplication-associated SNPs are unevenly distributed across chromosomes but coding and non-coding SNPs are affected equally

The genomic distribution of DI-genotypes revealed that CN3- and CN4-associated SNPs frequently occurred in runs of consecutive SNPs sharing the same dosage category (Fig. 6), as expected if these signatures originate from duplicated genomic segments. Runs ranged from a few hundred base pairs to several tens of kilobases, with similar length distributions for CN3 and CN4 states (Supplementary Figure 13).

**Figure 6.**
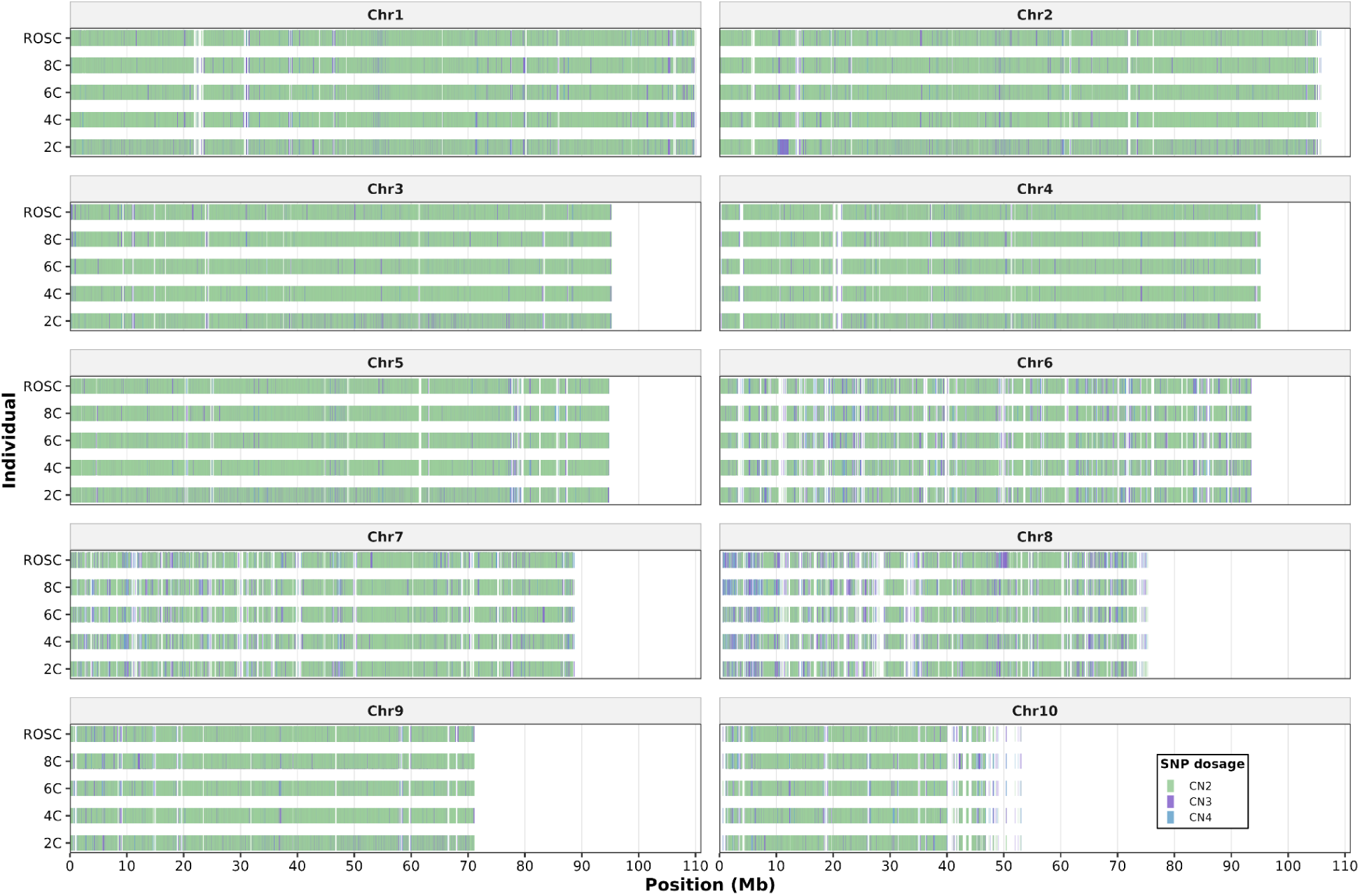
Density of duplication-associated SNPs. Plots are organized by chromosome for each individual. SNP dosage categories are indicated by colours (green = CN2, purple = CN3 and blue = CN4).

Although duplication-associated SNPs were detected on all chromosomes, their density differed markedly among chromosomes. Chromosomes 6, 7 and 8 consistently exhibited the highest densities of CN3 and CN4 SNPs in all five individuals (Figs. 6 and 7A). Moreover, these three chromosomes displayed substantially higher CN4/CN3 ratios (0.44, 0.36 and 0.42, respectively) than the remaining chromosomes (mean = 0.24), in accordance with duplicated (2-copy) segment alleles segregating at higher frequencies on these chromosomes.

**Figure 7.**
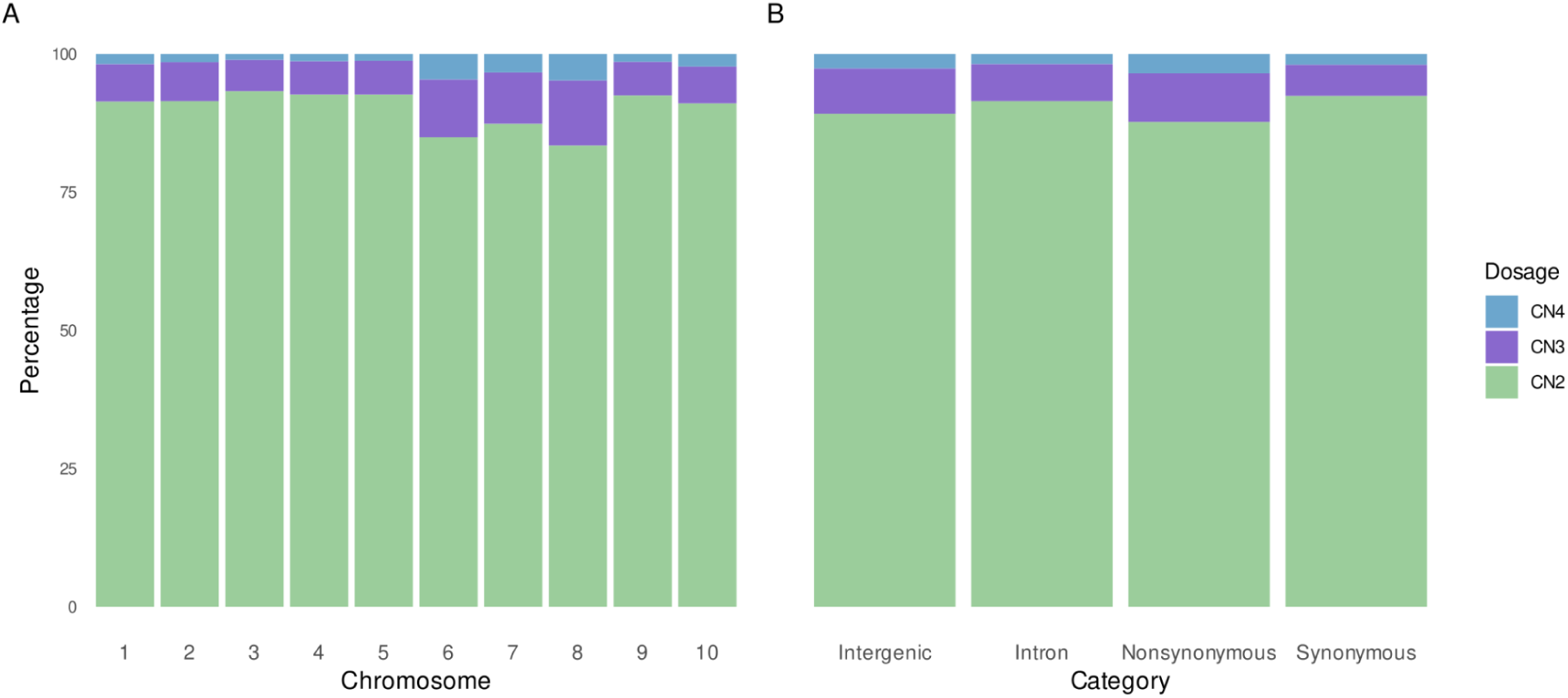
**A** Percentage of SNPs with a dosage of CN2, CN3 and CN4 in all 10 chromosomes, averaged across all five individuals and **B** percentage of intergenic, intronic, nonsynonymous and synonymous variants with a dosage of CN2, CN3 and CN4, averaged across all five individuals.

This chromosomal heterogeneity prompted us to examine what might distinguish chromosomes 6, 7 and 8 from the others. *O. edulis* is known to exhibit substantial somatic aneuploidy, a phenomenon commonly observed across bivalves, with chromosome losses strongly biased towards particular chromosomes (Thiriot-Quiévreux 2002; de Sousa et al. 2012; Khatir et al. 2022; Sullivan 2024). Interestingly, several lines of evidence converge to suggest that chromosomes 6, 7 and 8, which carry the highest densities of duplication-associated SNPs, are also preferentially lost in somatic cells. Cytogenetic studies have shown substantial variation in chromosome morphology within Ostreinae, with *O. edulis* characterized by five metacentric and five non-metacentric chromosome pairs (Thiriot-Quiévreux 1984; Li and Havenhand 1997; Leitão et al. 2002). Consistently, the first five chromosomes each display a pronounced central valley of nucleotide diversity, likely corresponding to a centromeric region, whereas no such valleys are observed in the central part of chromosomes 6 to 10 (Supplementary Figure 14). Importantly, somatic chromosome loss in *O. edulis* has repeatedly been reported specifically for chromosomes 6 to 10 (Thiriot-Quiévreux 1986). Thus, among the five non-metacentric chromosomes, the three showing the highest densities of duplication-associated SNPs are also those most likely to be preferentially lost through somatic aneuploidy.

We do not believe that somatic aneuploidy itself directly generates the CN3 and CN4 signatures observed here. Chromosome loss affects only a fraction of cells within an individual and would therefore be expected to produce intermediate allelic dosages and reduced sequencing depth below CN2 rather than the well-defined CN3 and CN4 clusters recovered by both Smudgeplot and DI-genotype analyses. Nevertheless, SNPs that could not be confidently assigned to any dosage category occurred significantly more frequently on chromosomes 6 to 8 (χ² tests, P < 2.2 × 10⁻¹⁶ for all individuals; Supplementary Statistics), consistent with somatic aneuploidy contributing to reduced relative depth and altered allelic coverage fractions on these chromosomes. Thus, while somatic aneuploidy may explain part of the unassigned signal, it does not account for the excess of discrete CN3 and CN4 states. The association between duplication-associated SNPs and chromosomes prone to somatic aneuploidy therefore points instead to two non-mutually exclusive evolutionary explanations.

Under an architecture-driven hypothesis, both phenomena would arise from the same intrinsic chromosome properties. Segmental duplications frequently originate through unequal recombination or replication errors (Ohta 2000; Zhang 2003), and such events may occur more readily in chromosomes prone to pairing or cohesion defects or enriched in repetitive sequences. Whole-genome alignments visualized with D-GENIES (Cabanettes and Klopp 2018) further indicate that chromosomes 6, 7, and 8 contain a higher density of repeated sequences than the remaining chromosomes (Supplementary Figure 15). Under this hypothesis, the same chromosome architecture that promotes segregation instability during mitosis would also increase the mutation rate of segmental duplications, explaining the association between somatic aneuploidy and duplication density without requiring a causal effect of one on the other.

Alternatively, under a compensation-driven hypothesis, recurrent somatic chromosome loss could create selection on duplicated segments containing genes from unstable chromosomes. Somatic chromosome loss exposes recessive deleterious mutations to negative selection and creates gene-dosage imbalance in affected cells (Sheltzer and Amon 2011; Gilchrist and Stelkens 2019; Vande Zande et al. 2023). Segmental duplications may buffer these effects by providing additional gene copies, thereby compensating for reduced dosage and restoring functional redundancy (Kondrashov 2012; Katju and Bergthorsson 2013; Hose et al. 2015; Basilicata and Valsecchi 2021). In this scenario, duplicated copies of genes located on chromosomes prone to somatic loss would preferentially accumulate or be maintained, irrespective of the genomic location of the additional copy. Interchromosomal duplications could indeed provide particularly effective buffering when the additional copy lies on a more stable chromosome.

Comparisons with the Pacific oyster provide additional support for an association between chromosome-specific instability and duplication. Qi et al. (2021, 2023) reported elevated densities of segmental duplications and copy-number variable regions on a subset of *Magallana gigas* (and *M. angulata*) chromosomes. Synteny analysis (Li et al. 2023) indicates that these chromosomes correspond primarily to chromosomes 6 to 10 in *O. edulis* (Supplementary Table 3). Independently, Bouilly et al. (2005) showed that somatic chromosome loss in *M. gigas* is concentrated on a subset of chromosomes, with the largest chromosome displaying the highest frequency of aneuploid cells. Based on chromosome size and synteny, chromosome 1 in *M. gigas*, by far the largest chromosome in this species and the most enriched in repeat content (Qi et al. 2021, 2023), most likely corresponds to chromosome 8 in *O. edulis*, one of the three chromosomes showing the highest density of duplication-associated SNPs here. Although these comparisons remain indirect, they suggest that chromosome-specific somatic instability and polymorphic segmental duplications may be linked across oysters, highlighting the need to identify these particular chromosomes across oyster species and to investigate their genomic and evolutionary dynamics separately from the rest of the genome.

Finally, we examined whether duplication-associated SNPs were preferentially associated with particular functional categories. CN2, CN3, and CN4 SNPs were distributed similarly among intergenic, intronic, synonymous, and non-synonymous sites (Fig. 7B). Although intergenic and non-synonymous sites showed a slightly lower proportion of CN2 SNPs than intronic and synonymous sites (χ² tests, P < 2.2 × 10⁻¹⁶ for all individuals; Supplementary Statistics), the differences were modest, indicating that duplicated genomic segments affect coding and non-coding regions to a similar extent. Functional enrichment analysis revealed no significant KEGG pathway after false discovery rate (FDR) correction for chromosomes 6 and 7, whereas five pathways remained significant for chromosome 8 (Supplementary Tables 4–6). Lysine degradation was among the strongest enrichments for both chromosomes 7 and 8, with 43 of the 61 genes assigned to this pathway located on these two chromosomes. Chromosome 8 was also enriched for N-glycan and glycosphingolipid biosynthesis pathways. Although these results should be interpreted cautiously, the latter pathways are involved in cell-surface interactions and immune functions, suggesting that immunity may warrant further investigation in relation to the accumulation or maintenance of duplications on these chromosomes. Overall, however, polymorphic duplications were broadly distributed across functional categories rather than preferentially associated with particular classes of genes, consistent with observations in *Crassostrea virginica* (Modak et al. 2021).

### CONCLUSIONS AND IMPLICATIONS FOR POPULATION GENETICS STUDIES

The high proportion of duplication-associated SNPs in *O. edulis*, independently supported by the het-mer and reference-based analyses, is consistent with the extensive structural variation previously reported in other oysters (e.g. Modak et al. 2021; Qi, Li, and Zhang 2021). However, the magnitude of the allelic-coverage distortion within individuals, with approximately 30% of heterozygous SNP calls having an allelic coverage fraction between 0.2 and 0.4, remains a surprise to us. This raises two broader questions: First, how should low or moderate coverage datasets be analyzed for population structure, demographic reconstruction, and selection scans in such genomes? And what evolutionary processes generate and maintain such extensive polymorphic duplication in oysters, and perhaps more generally in marine bivalves?

The availability of five high-coverage genomes allowed us to examine structural variation and its impact on SNP calling in a way that is rarely possible in population genomic datasets. Typically, only the reference individual is deeply sequenced, whereas resequenced individuals have insufficient depth for the het-mer and dosage-based inference used here. Short indels and some CNVs can be called from short reads, but large or recent duplications remain difficult to resolve even with paired-end, linked-read, or long-read approaches when copies are long, highly similar, and possibly inter-chromosomal. High-coverage dosage inference therefore provides useful complementary information. By contrast, filtering a low or moderate coverage dataset using depth from a single individual is unlikely to be sufficient because the expected 2X, 3X, and 4X depth distributions and the corresponding allelic coverage fractions overlap substantially at low coverage, and polymorphic duplications occur at different loci in different individuals. Joint information across multiple individuals can partly alleviate this problem. Methods such as ngsParalog (Linderoth 2018) and ParaMask (Tjeng et al. 2025) use sample-wide information to identify deviant SNPs in multicopy regions, including heterozygote frequencies and the expectation that problematic SNPs cluster in duplicated segments. However, these methods were primarily designed for fixed or collapsed multicopy regions and may miss rare polymorphic duplications or CNVs that deviate little from Hardy-Weinberg expectations. We therefore recommend explicit inspection of allelic coverage and depth distributions and, when possible, filtering of duplication-associated SNPs. For *O. edulis*, exclusion of chromosomes 6, 7, and 8 may also be considered for analyses particularly sensitive to such variants, such as demographic inference or population structure. In other oyster species, chromosomes syntenic to these three chromosomes should similarly be identified and either excluded from sensitive analyses or, at minimum, examined separately for unusual patterns of coverage and structural variation. We provide a supplementary file listing CN3 and CN4 SNPs that can be directly filtered in studies using the same *O. edulis* reference genome.

Why *O. edulis*, Ostreoidea, and perhaps bivalves more generally, harbor such extensive polymorphic duplication remains unresolved. Pangenome studies will be needed to distinguish the mechanistic constraints and evolutionary forces shaping these patterns. Ancient whole-genome duplications proposed near the origin of Ostreidae or in *Mytilus* (Corrochano-Fraile et al. 2022) may have contributed to long-term copy-number dynamics but cannot by themselves explain the segregating duplications observed here. The duplicated copies detected by our approaches are likely to be weakly diverged, either because the duplications are recent enough to remain polymorphic or because highly diverged copies would no longer collapse or co-map efficiently. Consistent with their predominantly low inferred frequencies, the reference genome most often appears to carry the 1-copy segment allele, or occasionally a collapsed representation of the duplicated state. Although larger population samples will be required to formally test the frequency spectrum of 2-copy segment alleles against neutral expectations, the observed frequency spectrum and the similar proportions of singleton 2-copy segment and standard CN2 SNP alleles in our small sample are consistent with polymorphic duplications being largely neutral or nearly neutral (i.e. slightly deleterious). The high burden of polymorphic duplications may therefore largely reflect the exceptionally high genetic diversity and large population sizes of oysters, allowing numerous neutral or slightly deleterious structural variants to segregate. The broadly similar representation of duplication-associated SNPs at non-coding, synonymous and non-synonymous sites is also compatible with predominantly weak selective effects. The association between duplication-associated SNPs and chromosomes prone to somatic aneuploidy nevertheless suggests a possible effect of selection on their chromosomal distribution. If some duplications partially buffer the consequences of somatic chromosome loss, as proposed by our compensation-driven hypothesis, their selective advantage at the individual level is likely to be extremely small. Indeed, the persistence of oysters with substantial variation in somatic aneuploidy within natural populations suggests that its effects on individual fitness are relatively modest, and therefore that selection for mechanisms buffering these effects should also be weak. Such weak selection would be compatible with the nearly neutral frequency patterns observed here, while potentially influencing the long-term accumulation of duplications associated with chromosomes prone to somatic instability. The architecture-driven hypothesis, in contrast, does not require selection. Chromosome-specific differences in duplication mutation rates could generate the observed heterogeneity even if most duplications are neutral or nearly neutral. Stronger adaptive and maladaptive effects may nevertheless also contribute. The possible role of presence-absence and duplication variation in the evolutionary success, phenotypic variation, and local adaptation of marine bivalves has been emphasized by others (e.g. Gerdol et al. 2020; Modak et al. 2021). We suggest instead that the high genetic load of these abundant, highly fecund organisms with sweepstakes reproduction (Hedgecock and Pudovkin 2011; Harrang et al. 2013; Plough 2016) may provide a simpler explanation for the segregation of numerous mildly deleterious copy-number variants, paralleling the high burden of mildly deleterious non-synonymous variation reported in *O. edulis* (Harrang et al. 2013). Population-genetic analyses based on accurately resolved duplication variants will be required to distinguish among these alternatives.

## Acknowledgements

The authors are very grateful to Antonio Villalba for carrying out the sampling of the four Galician flat oyster samples used in this study. They also express their thanks to Xavier Dallaire, Pierre-Alexandre Gagnaire, Céline Reisser, and Christelle Fraïsse for valued discussion and insight.

## Data, scripts, code and supplementary information availability

The four newly sequenced *O. edulis* whole genomic read datasets are available at: https://www.ebi.ac.uk/ena/browser/view/PRJEB93918

Code used in this study is available at: https://doi.org/10.5281/zenodo.21297519

A SNP list of duplicated regions generated from high coverage individuals is available at: https://zenodo.org/records/21510817.

Supplementary figures, tables and statistics are available after references below.

## SUPPLEMENTARY TABLES

**Supplementary Table 1.**
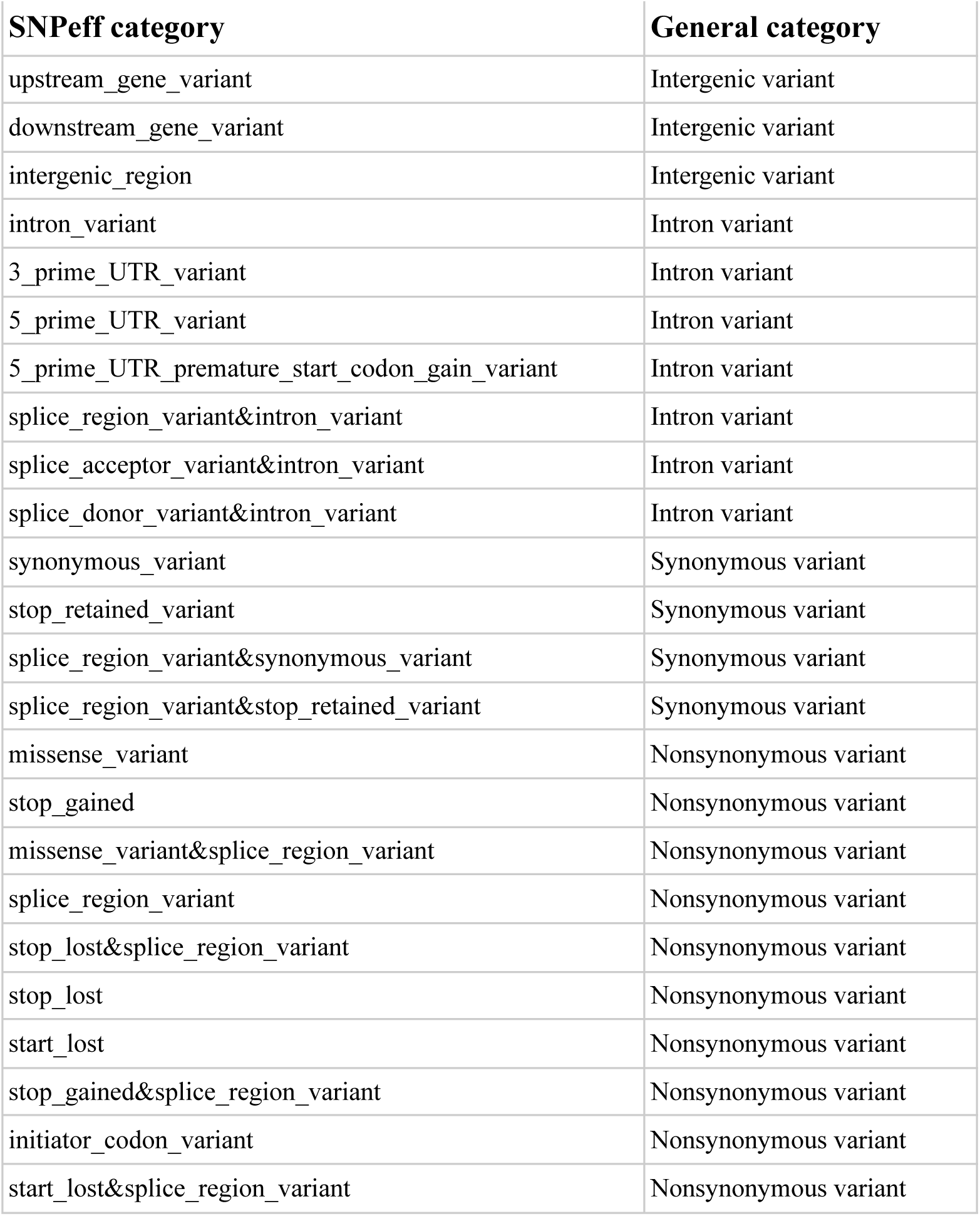
Variant categories assigned by SNPeff and the general category assigned for use in this study.

**Supplementary Table 2.**
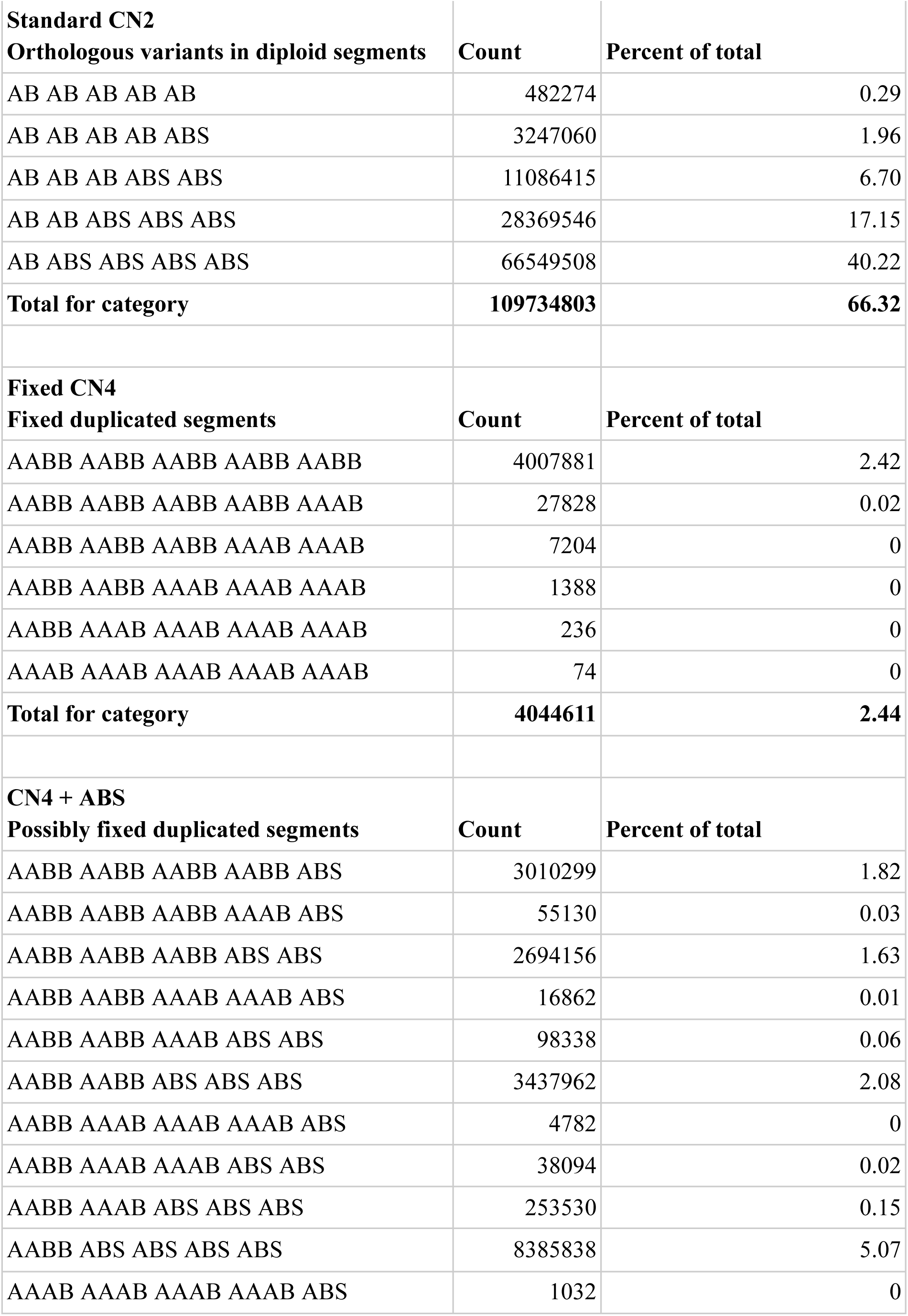
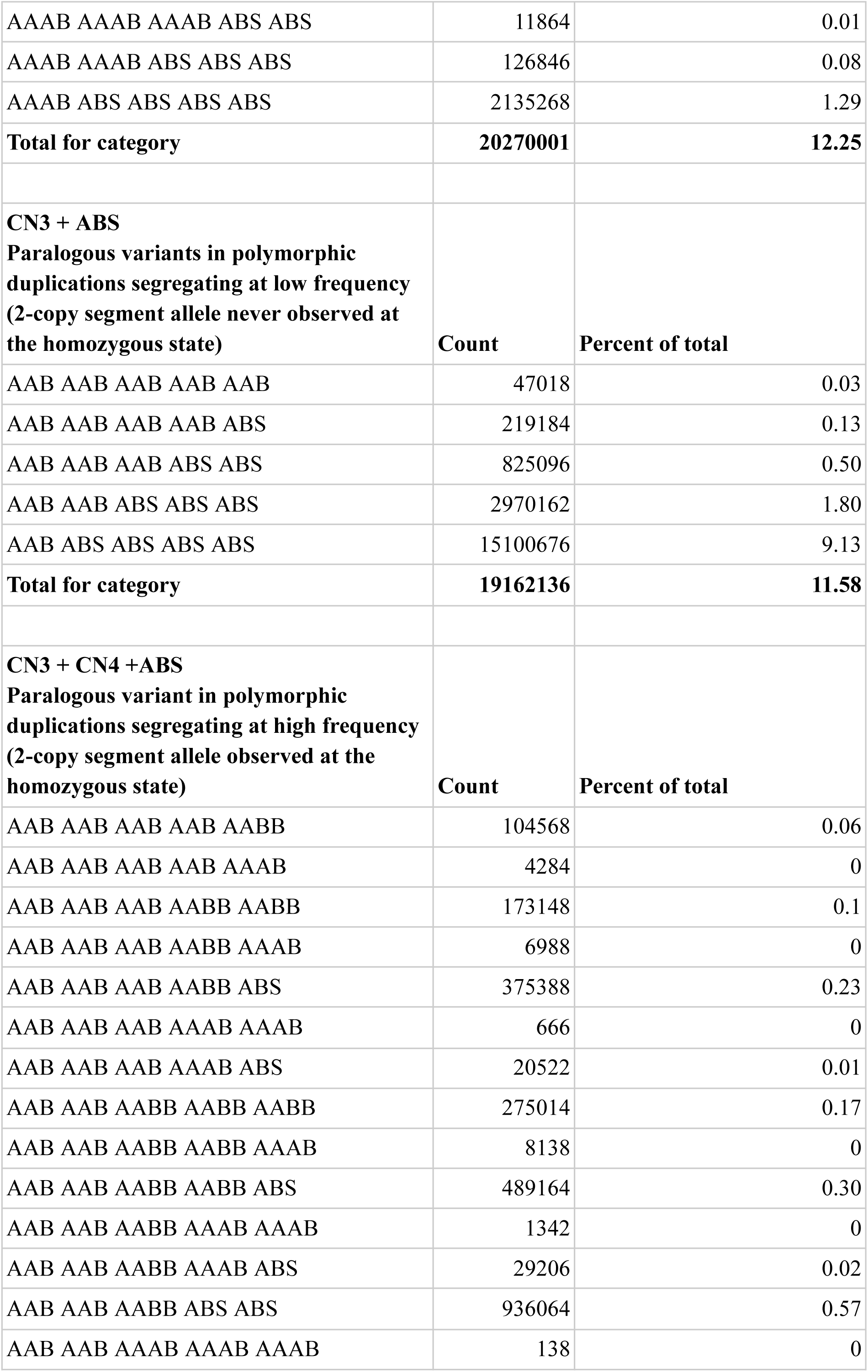
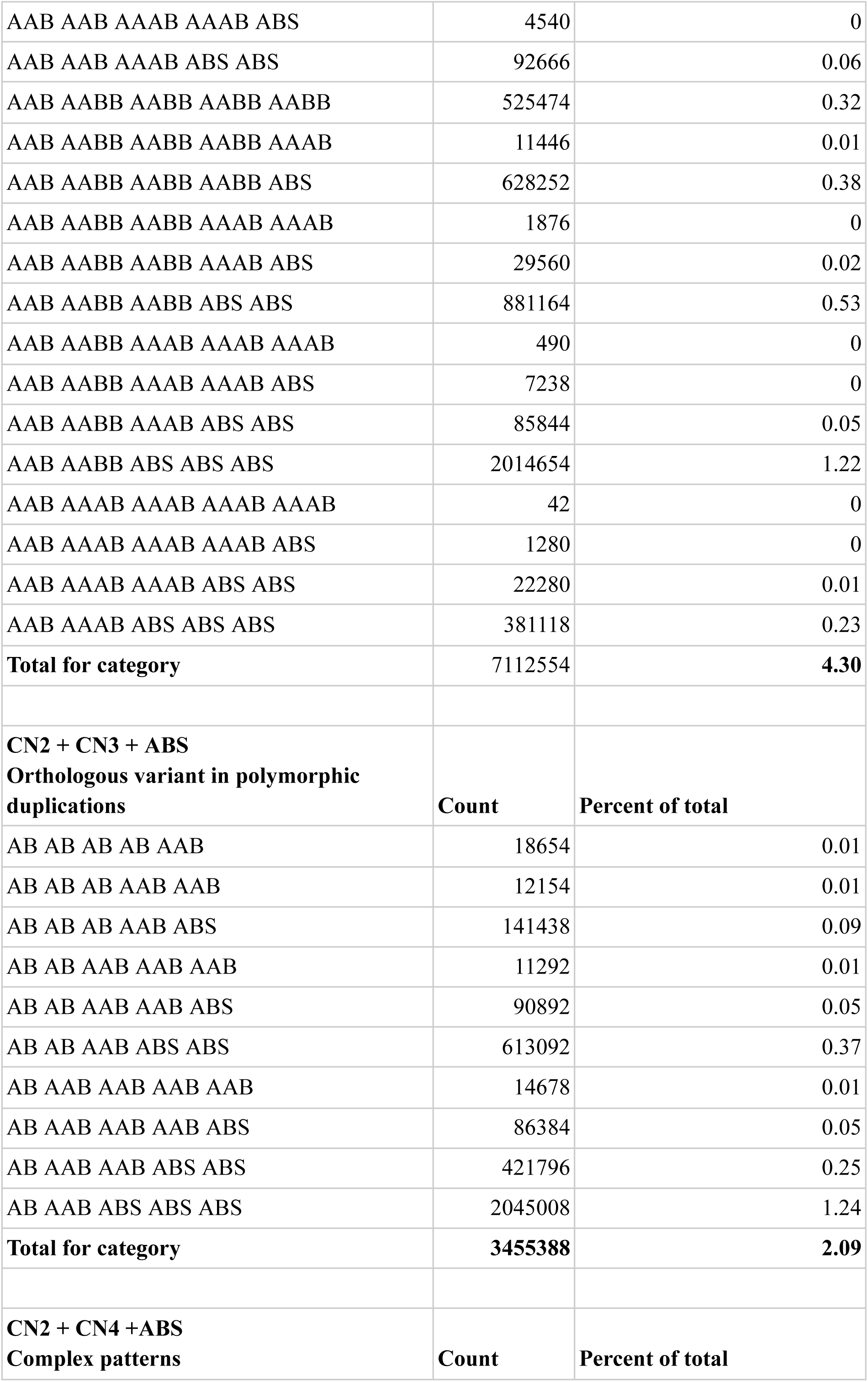
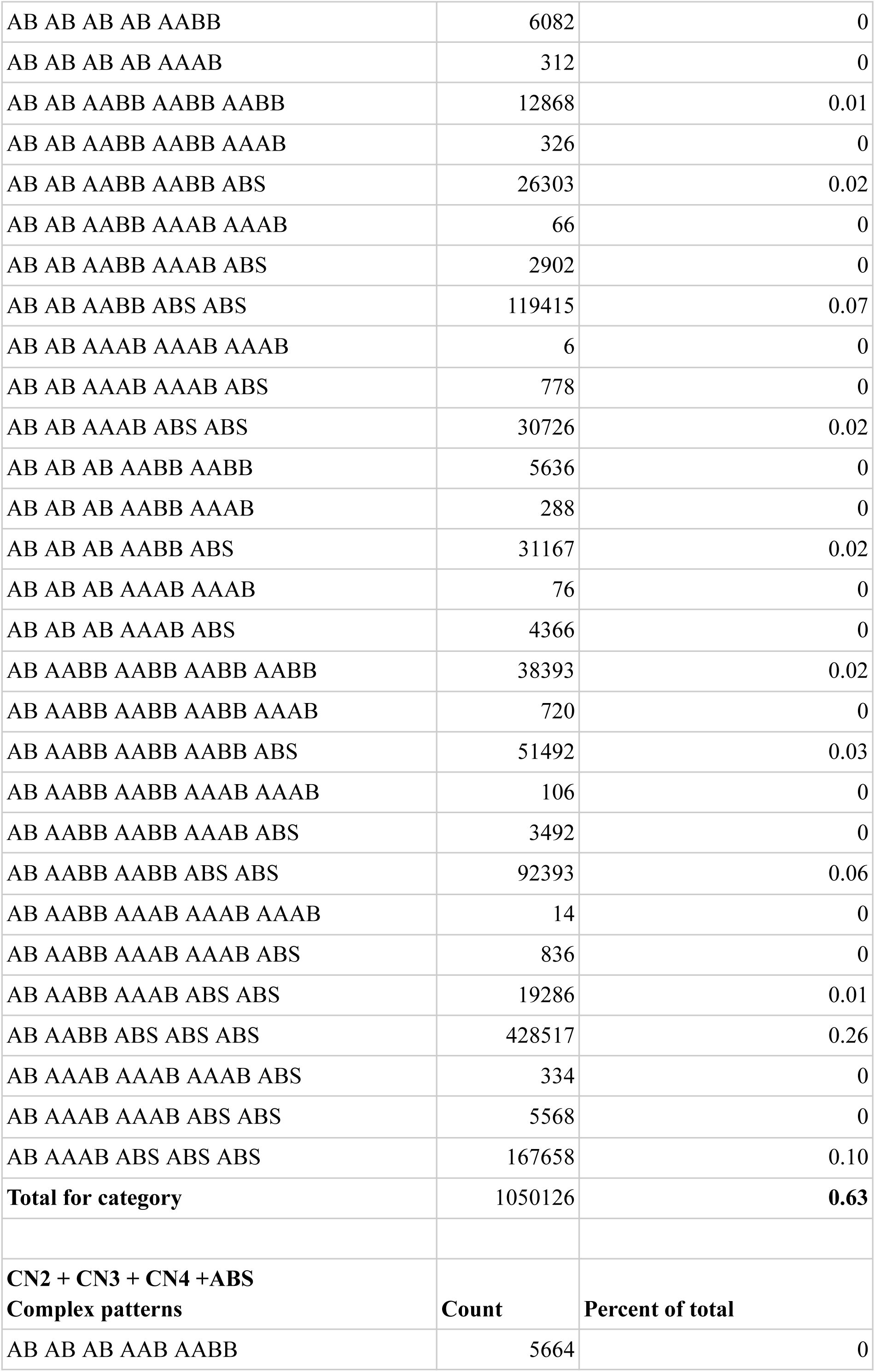
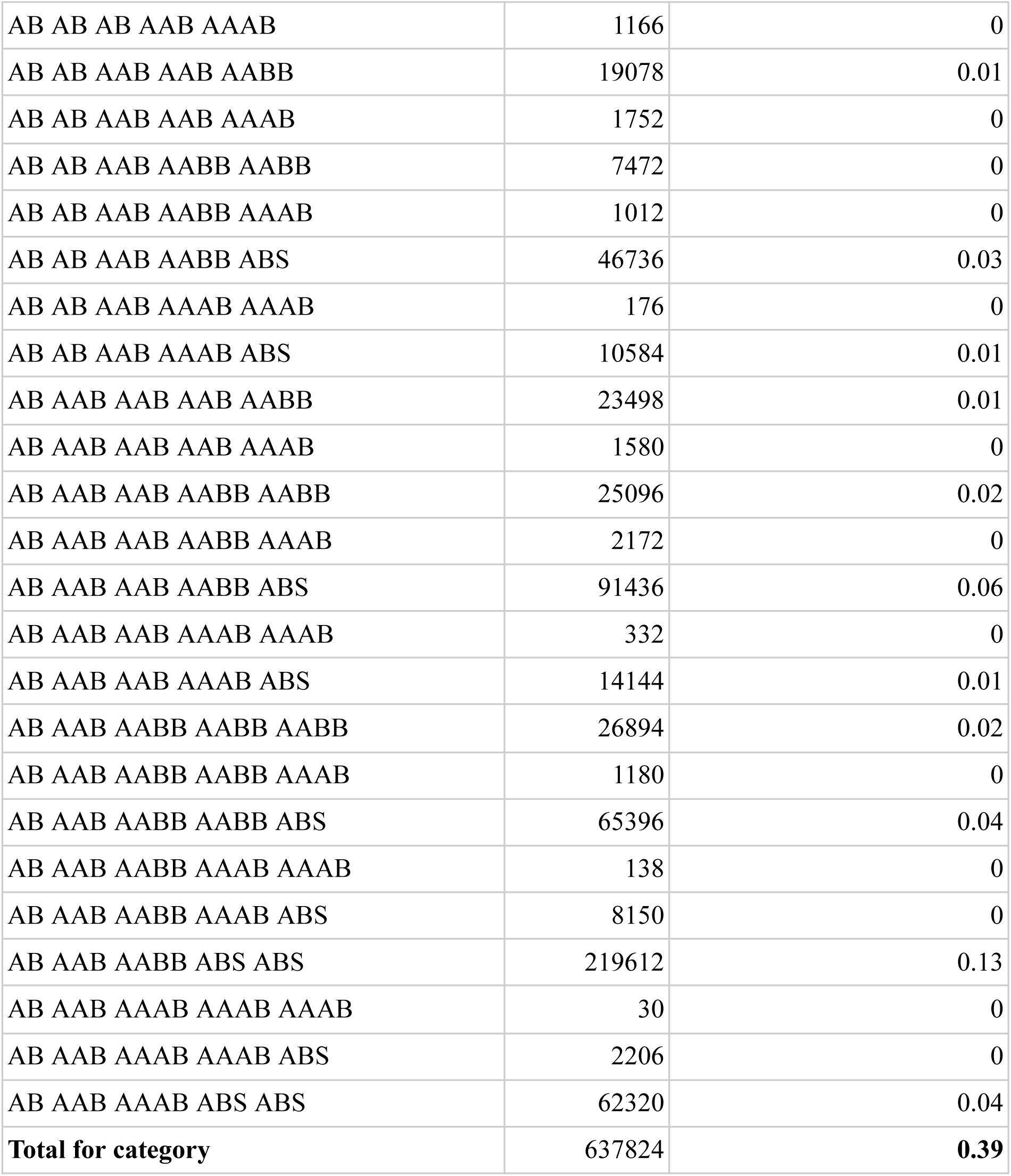
Comparison table containing the count of instances that a het-mer marker had each possible combination of genotypes identified in the five individuals. Subcategories organized for the different types of combination that were observed.

**Supplementary Table 3.** Correspondence between chromosomes as numbered in this study and *M. gigas* chromosomes as numbered by Qi, Li, and Zhang (2021) based on synteny analysis by Li et al. (2023), and corresponding chromosome ordering of alternate *O. edulis* reference genomes. Note that the reference genome by Boutet et al. (2022) was used by Lapègue et al. (2023) to identify putative inversions on chromosome numbers marked with an asterisk. These putative inversions were confirmed by Sambade et al. (2022) and Alves Monteiro et al. (2024).

| *O. edulis* (Li et al. 2023, **this study**) | *M. gigas* (Qi, Li, and Zhang 2021) | *O. edulis* (Boutet et al. 2022) | *O. edulis* (Gundappa et al. 2022) | *O. edulis* (Adkins and Mrowicki 2023) |
| --- | --- | --- | --- | --- |
| **1** | 2 | 1 | 1 | 1 |
| **2** | 5 | 2 | 4 | 2 |
| **3** | 9 | 3 | 6 | 5 |
| **4** | 7 | 4* | 5 | 6 |
| **5** | 6 | 5* | 2 | 4 |
| **6** | 3 | 6 | 3 | 3 |
| **7** | 4 | 7 | 7 | 7 |
| **8** | 1 | 9 | 9 | 8 |
| **9** | 8 | 8* | 8 | 9 |
| **10** | 10 | 10 | 10 | 10 |

**Supplementary Table 4.** GO terms significantly enriched for chromosome 6 at p < 0.05. None are significant after FDR correction.

| GO Term | Database | ID | Input number | Background number | P-Value | Corrected P-Value |
| --- | --- | --- | --- | --- | --- | --- |
| Pentose and glucuronate interconversions | KEGG PATHWAY | crg00040 | 7 | 20 | 5.32E-03 | 5.38E-01 |
| Terpenoid backbone biosynthesis | KEGG PATHWAY | crg00900 | 6 | 21 | 2.08E-02 | 8.07E-01 |
| Neuroactive ligand-receptor interaction | KEGG PATHWAY | crg04080 | 36 | 276 | 2.69E-02 | 8.07E-01 |
| Glyoxylate and dicarboxylate metabolism | KEGG PATHWAY | crg00630 | 9 | 45 | 3.19E-02 | 8.07E-01 |
| Phenylalanine, tyrosine and tryptophan biosynthesis | KEGG PATHWAY | crg00400 | 3 | 7 | 4.36E-02 | 8.81E-01 |

**Supplementary Table 5.**
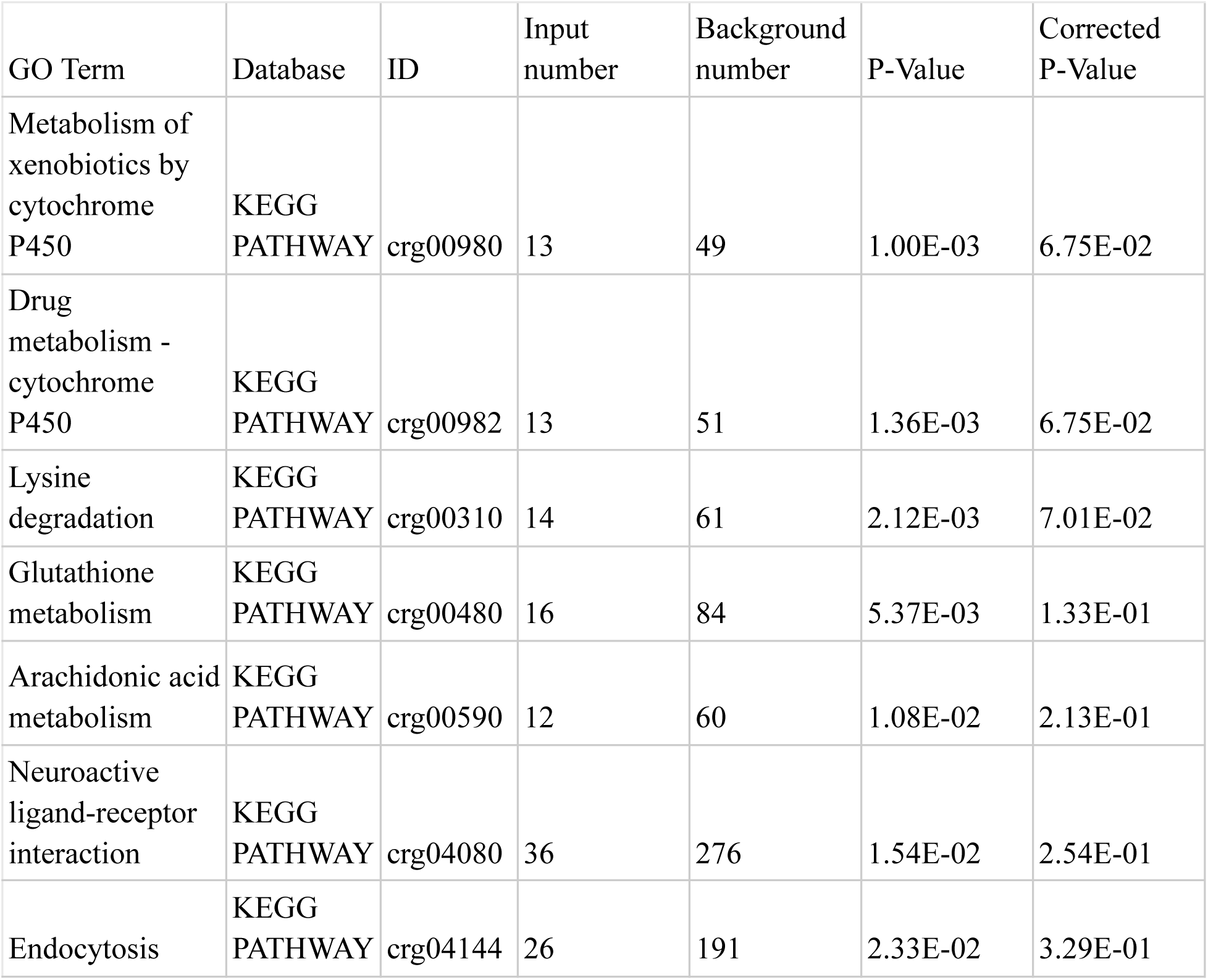
GO terms significantly enriched for chromosome 7 at p < 0.05. None are significant after FDR correction.

| GO Term | Database | ID | Input number | Background number | P-Value | Corrected P-Value |
| --- | --- | --- | --- | --- | --- | --- |
| Metabolism of xenobiotics by cytochrome P450 | KEGG PATHWAY | crg00980 | 13 | 49 | 1.00E-03 | 6.75E-02 |
| Drug metabolism - cytochrome P450 | KEGG PATHWAY | crg00982 | 13 | 51 | 1.36E-03 | 6.75E-02 |
| Lysine degradation | KEGG PATHWAY | crg00310 | 14 | 61 | 2.12E-03 | 7.01E-02 |
| Glutathione metabolism | KEGG PATHWAY | crg00480 | 16 | 84 | 5.37E-03 | 1.33E-01 |
| Arachidonic acid metabolism | KEGG PATHWAY | crg00590 | 12 | 60 | 1.08E-02 | 2.13E-01 |
| Neuroactive ligand-receptor interaction | KEGG PATHWAY | crg04080 | 36 | 276 | 1.54E-02 | 2.54E-01 |
| Endocytosis | KEGG PATHWAY | crg04144 | 26 | 191 | 2.33E-02 | 3.29E-01 |

**Supplementary Table 6.** GO terms significantly enriched for chromosome 8 at p < 0.05. Five are significant after FDR correction (indicated with a *).

| GO Term | Database | ID | Input number | Background number | P-Value | Corrected P-Value |
| --- | --- | --- | --- | --- | --- | --- |
| Lysine degradation* | KEGG PATHWAY | crg00310 | 29 | 61 | 2.96E-15 | 2.10E-13 |
| N-Glycan biosynthesis* | KEGG PATHWAY | crg00510 | 16 | 59 | 2.54E-06 | 9.01E-05 |
| Neuroactive ligand-receptor interaction* | KEGG PATHWAY | crg04080 | 31 | 276 | 8.66E-04 | 1.69E-02 |
| Taurine and hypotaurine metabolism* | KEGG PATHWAY | crg00430 | 6 | 16 | 9.51E-04 | 1.69E-02 |
| Glycosphingolipid biosynthesis-ganglio series* | KEGG PATHWAY | crg00604 | 6 | 20 | 2.43E-03 | 3.45E-02 |
| Glycosphingolipid biosynthesis-globo and isoglobo series | KEGG PATHWAY | crg00603 | 7 | 39 | 1.22E-02 | 1.44E-01 |
| Cysteine and methionine metabolism | KEGG PATHWAY | crg00270 | 8 | 53 | 1.81E-02 | 1.83E-01 |
| Glycosphingolipid biosynthesis-lacto and neolacto series | KEGG PATHWAY | crg00601 | 7 | 45 | 2.30E-02 | 2.04E-01 |
| TGF-beta signaling pathway | KEGG PATHWAY | crg04350 | 7 | 53 | 4.57E-02 | 3.32E-01 |
| Glycosaminoglycan degradation | KEGG PATHWAY | crg00531 | 6 | 42 | 4.67E-02 | 3.32E-01 |

## SUPPLEMENTARY FIGURES

**Supplementary Figure 1.**
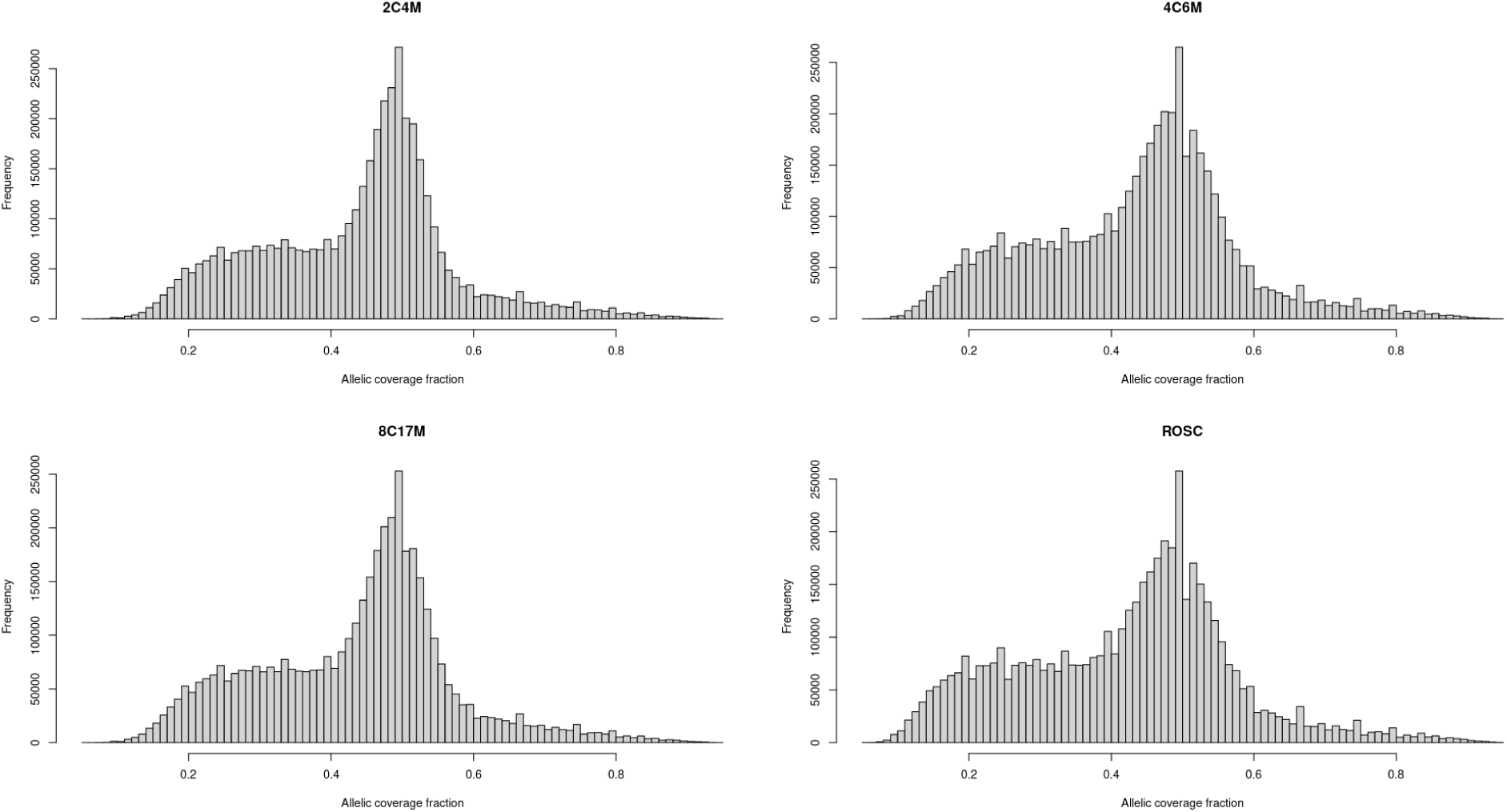
Histograms demonstrating the distribution of the allelic coverage fraction for called heterozygous sites in individual 2C4M, 4C6M, 8C17M and ROSC.

**Supplementary Figure 2.**
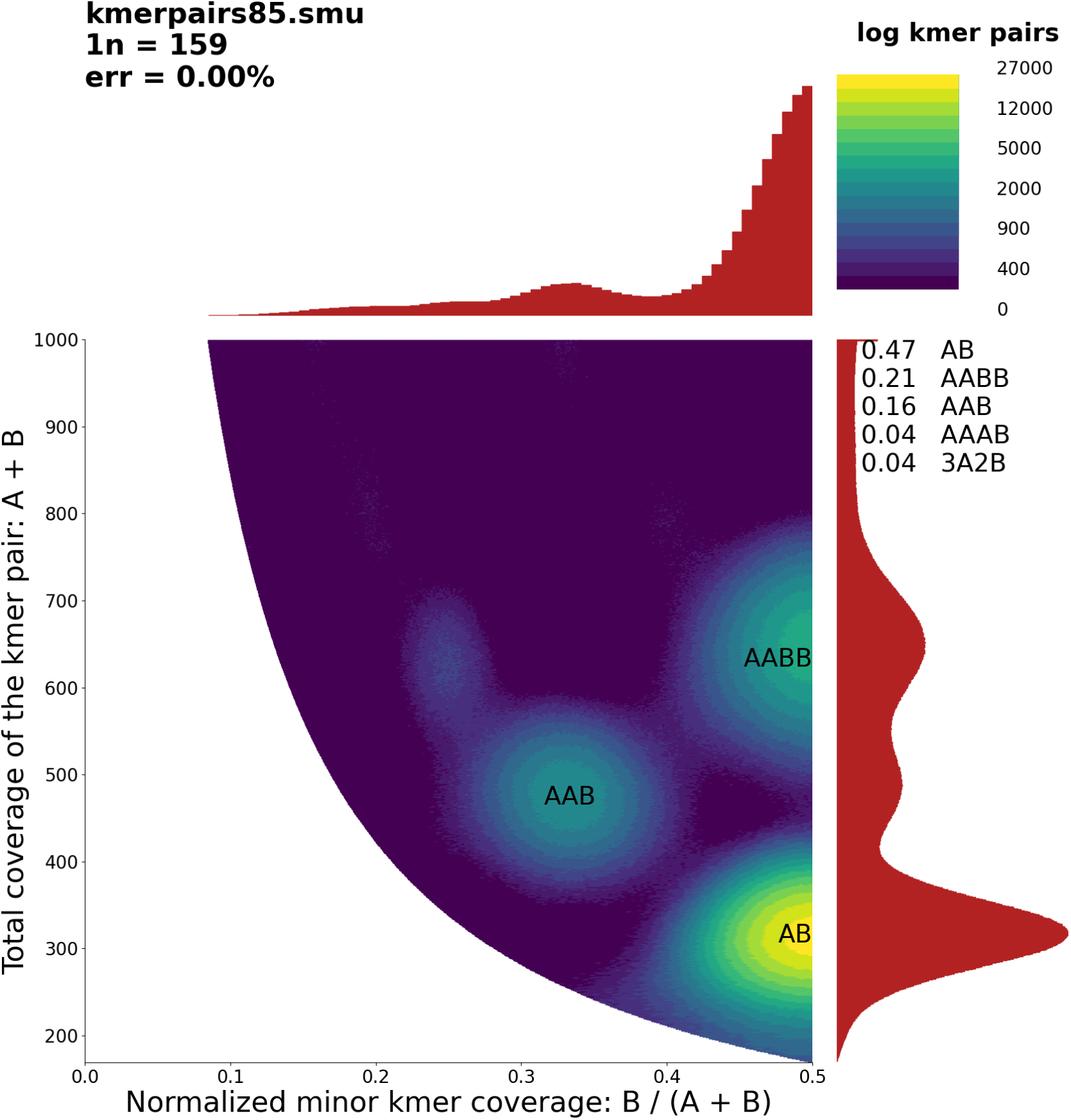
Smudgeplot analysis for sample 2C4M using an error threshold of 85 and a coverage range of 50-200.

**Supplementary Figure 3.**
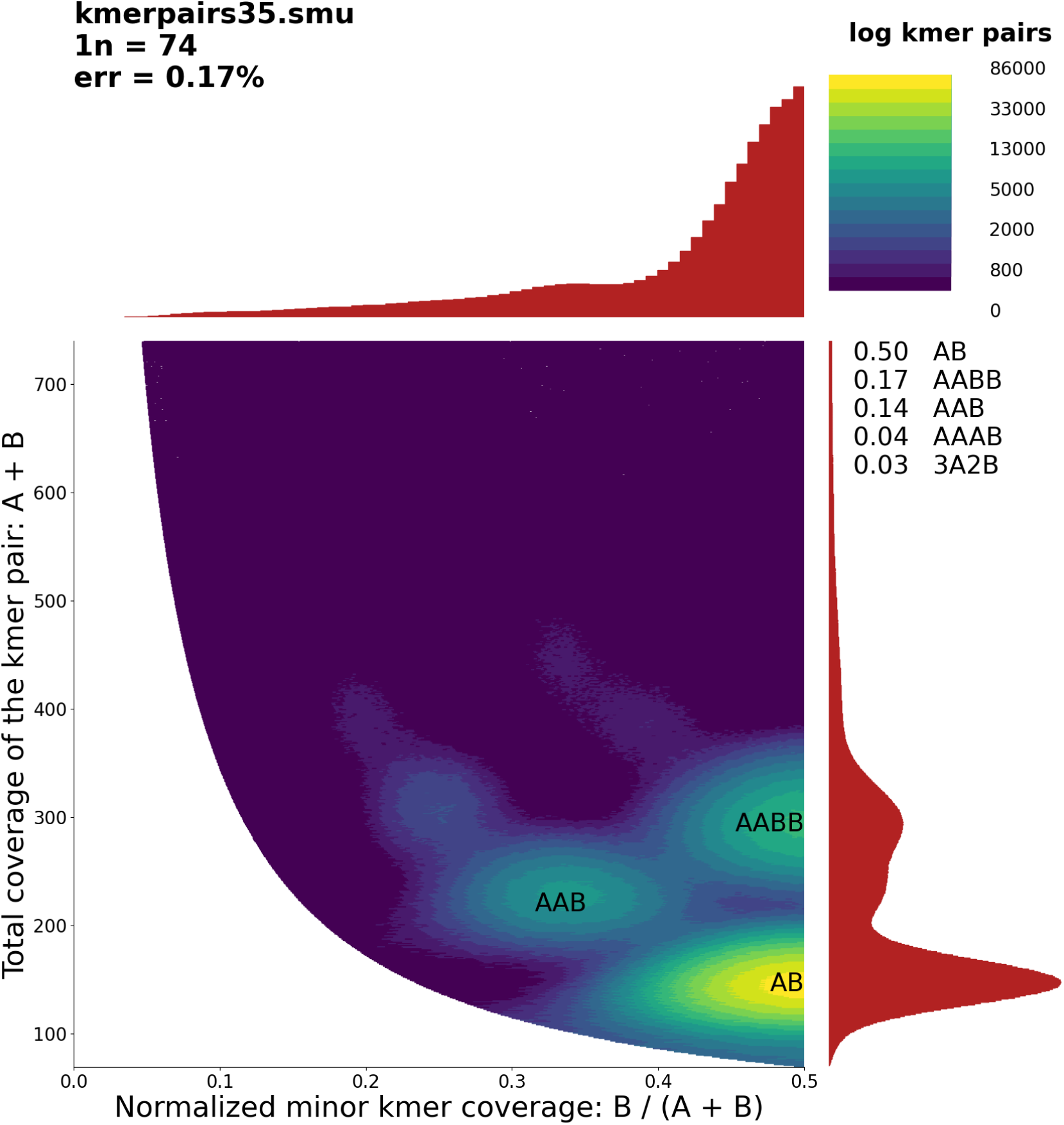
Smudgeplot analysis for sample 4C6M using an error threshold of 35 and a coverage range of 50-110.

**Supplementary Figure 4.**
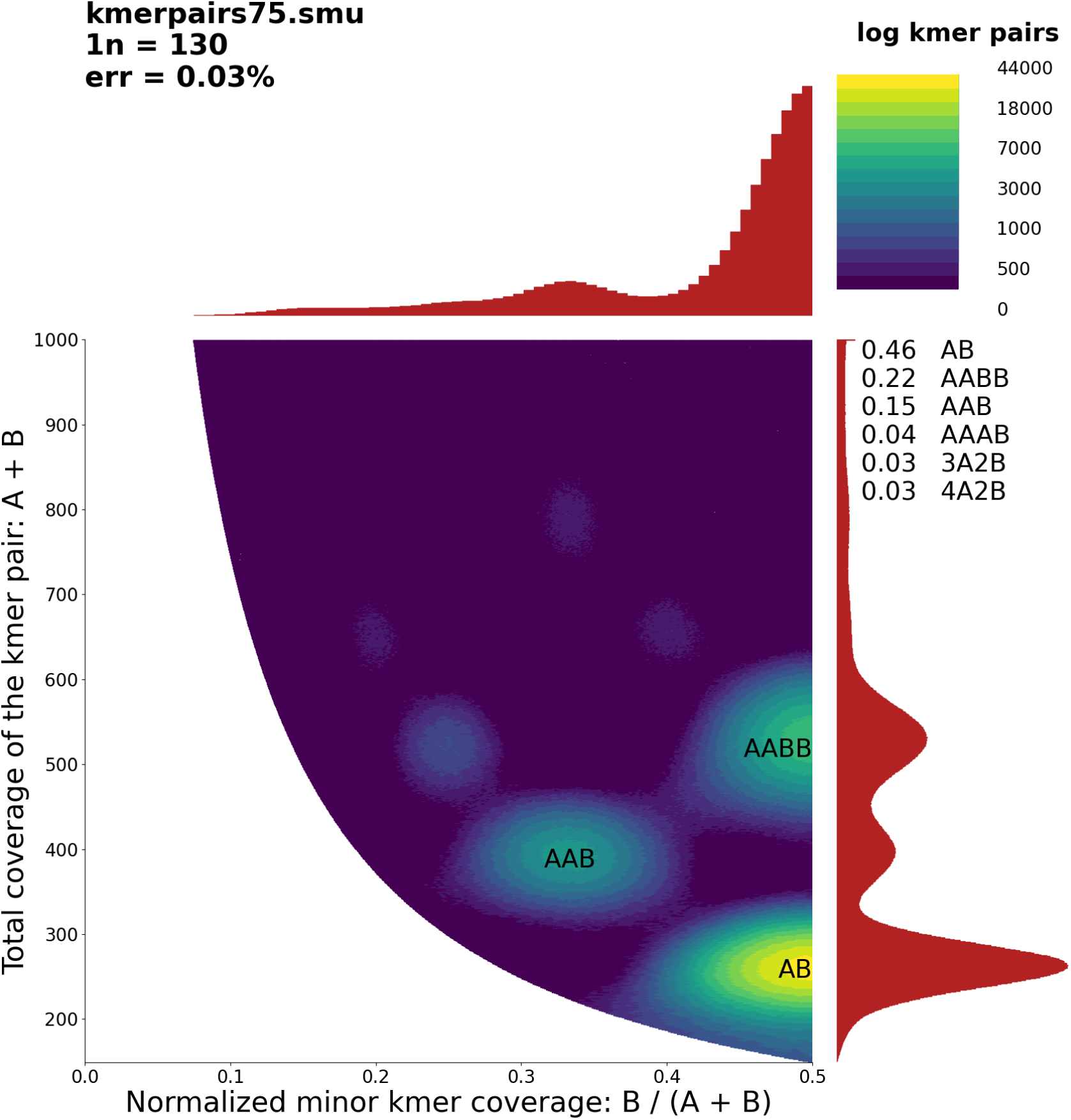
Smudgeplot analysis for sample 8C17M using an error threshold of 75 and a coverage range of 50-200.

**Supplementary Figure 5.**
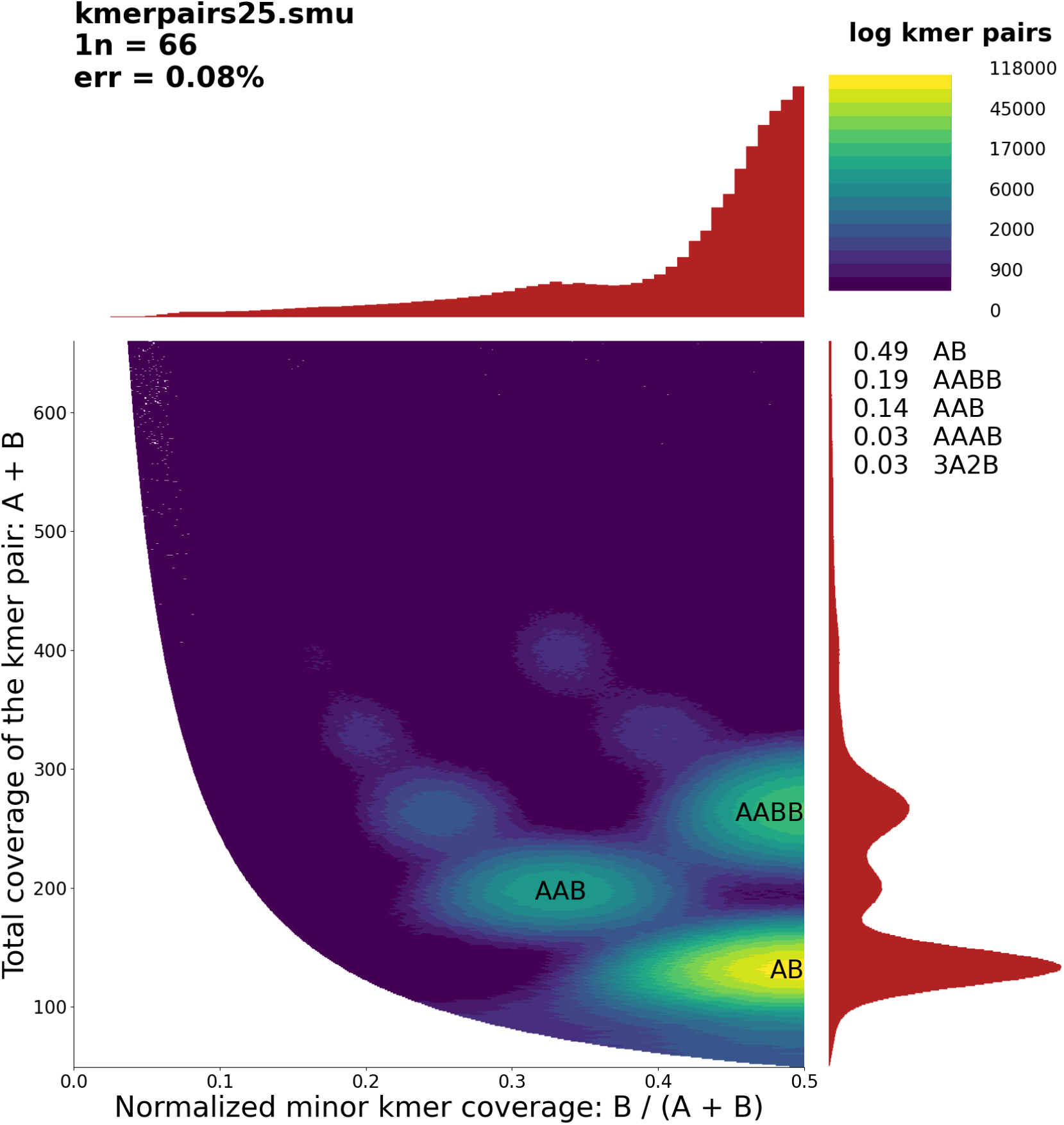
Smudgeplot analysis for sample ROSC using an error threshold of 25 and a coverage range of 40-100.

**Supplementary Figure 6.**
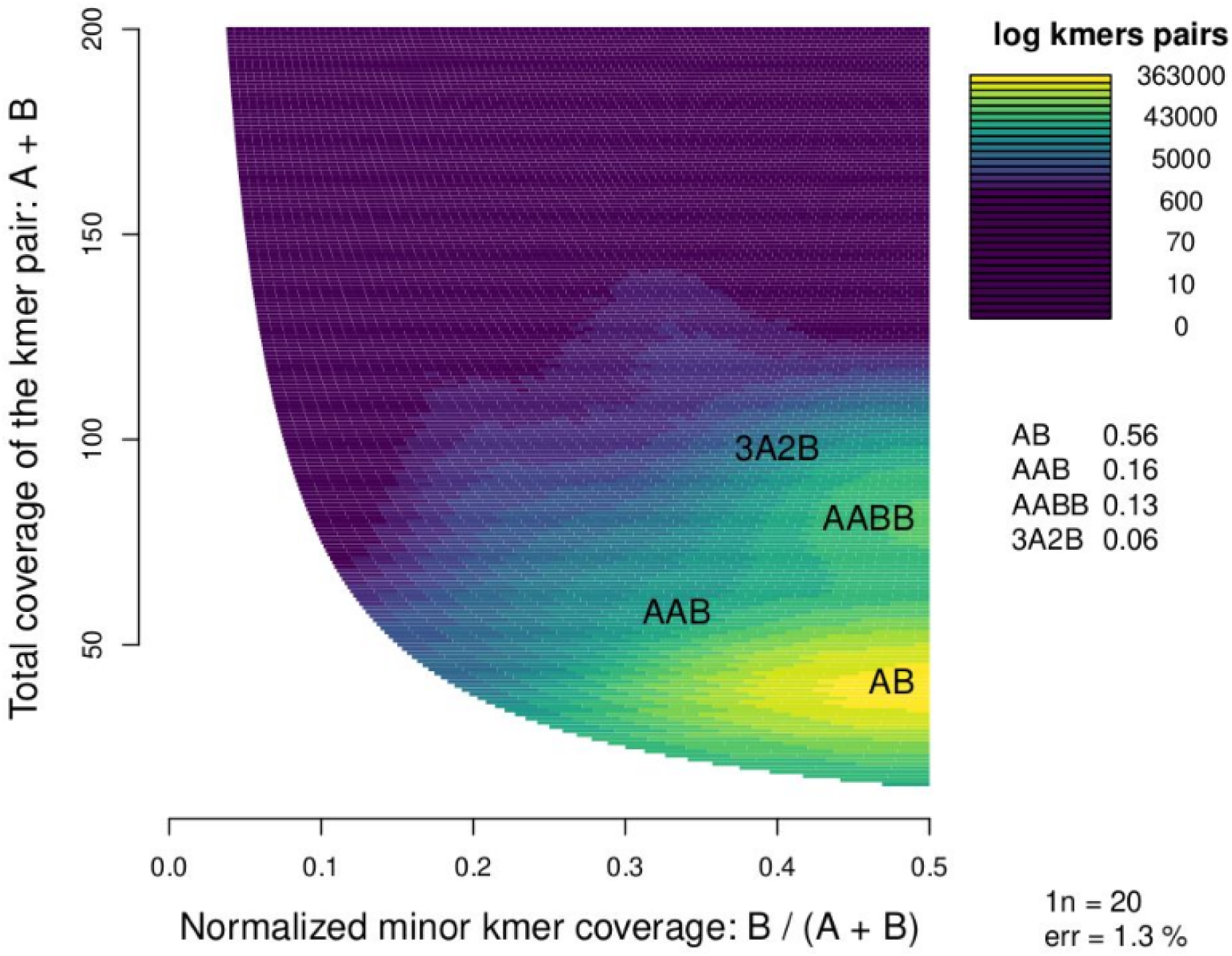
Smudgeplot analysis for *O. denselamellosa* using an error threshold of 8 and a coverage range of 5-50.

**Supplementary Figure 7.**
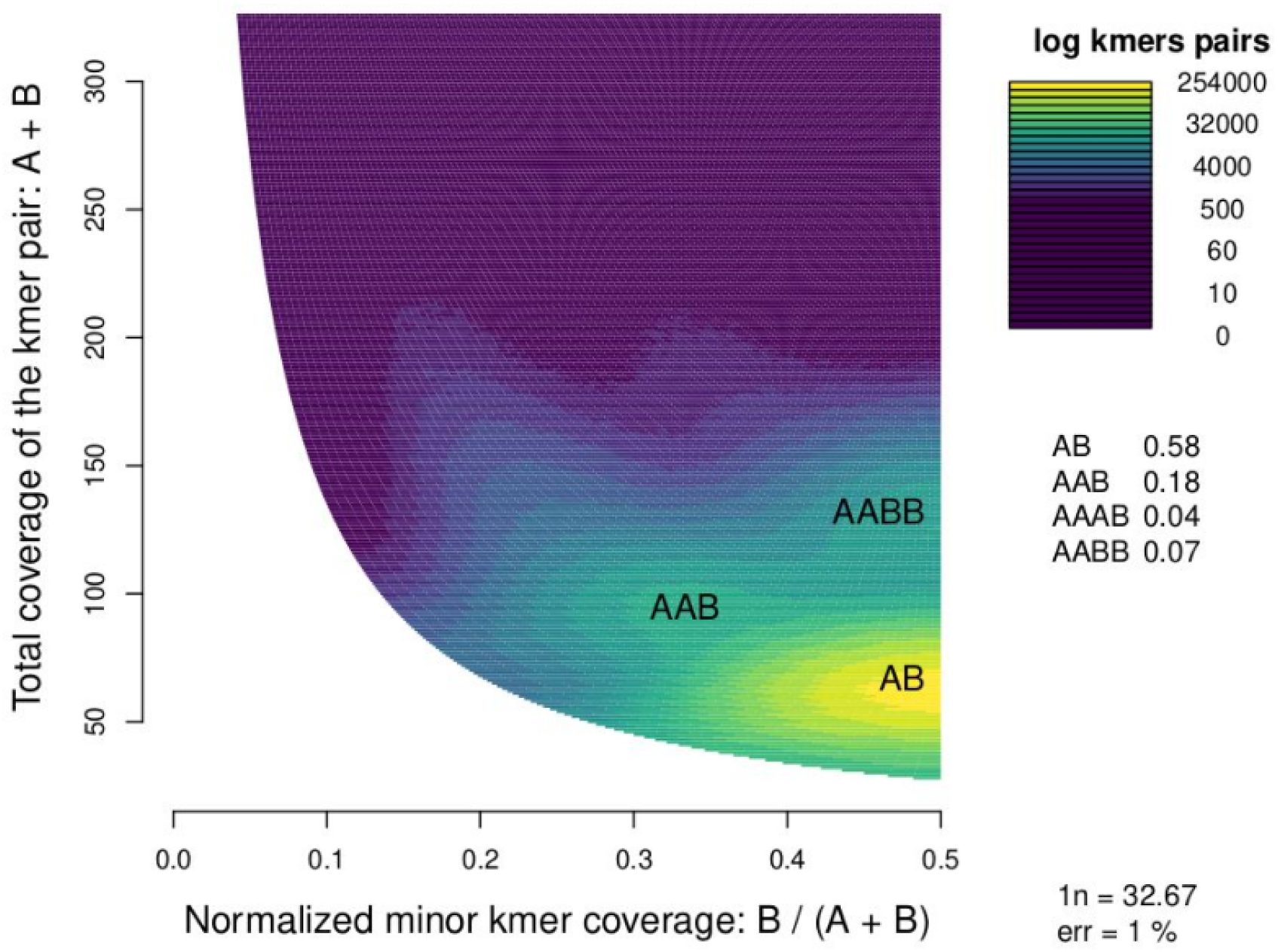
Smudgeplot analysis for *O. lurida* using an error threshold of 14 and a coverage range of 5-60.

**Supplementary Figure 8.**
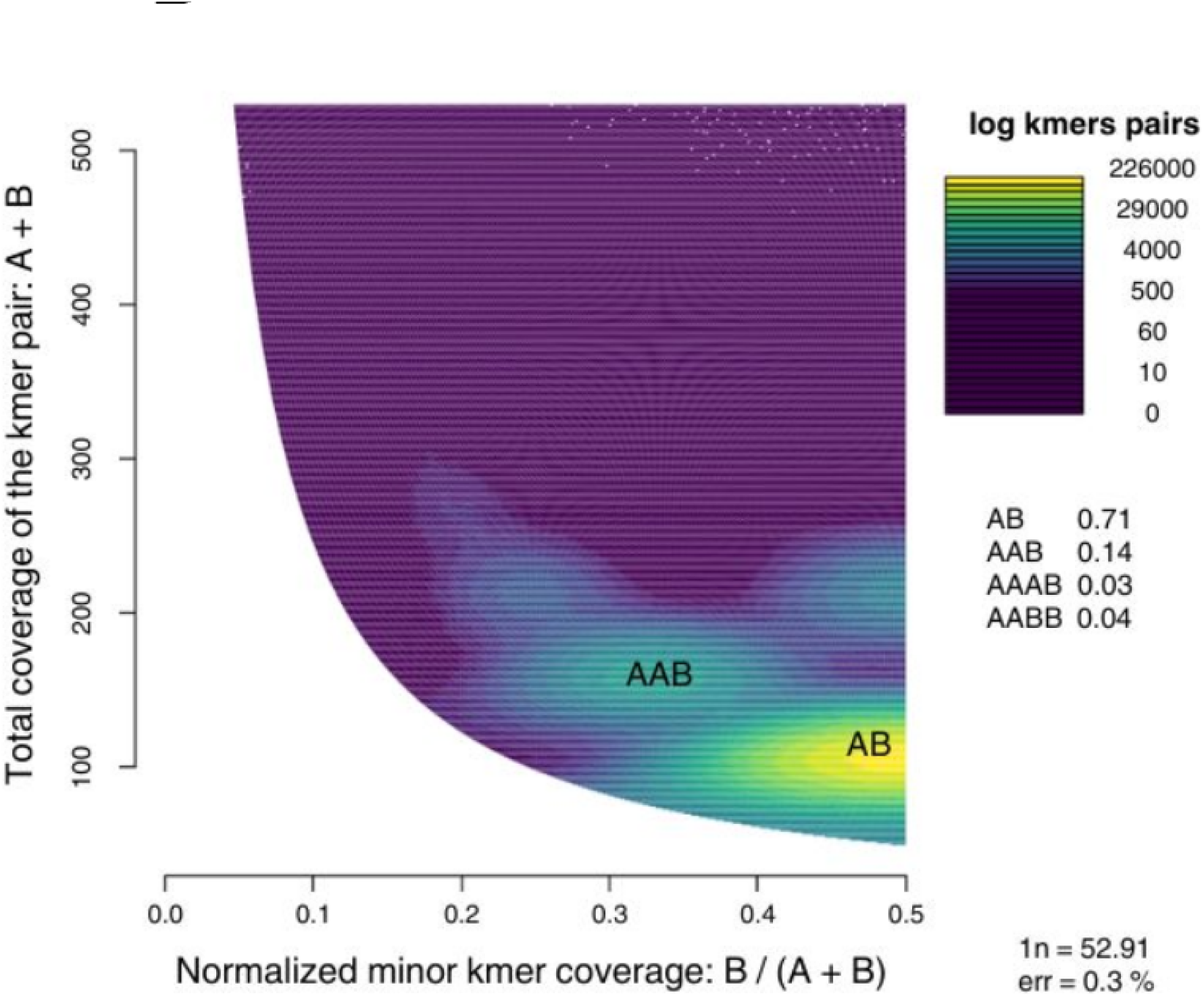
Smudgeplot analysis for *M. gigas* using an error threshold of 25 and a coverage range of 30-100.

**Supplementary Figure 9.**
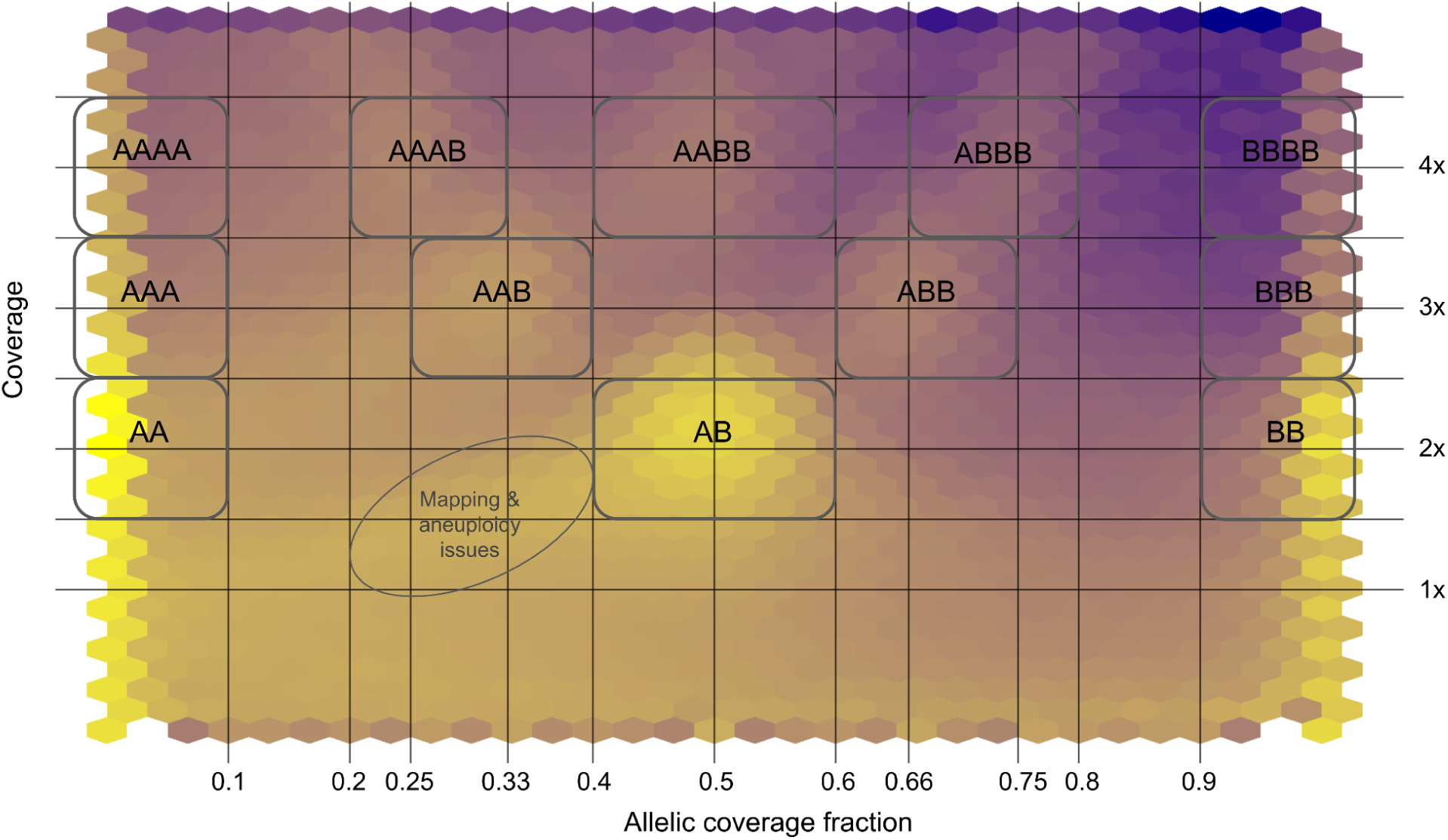
Density plot for sample 2C4M showing the joint distribution of allelic coverage fraction (x-axis) and relative sequencing depth (y-axis). Relative depth corresponds to SNP coverage normalized by the estimated haploid genome coverage, with the expected CN2, CN3 and CN4 depths indicated as 2X, 3X and 4X, respectively. Higher SNP densities are shown in brighter yellow. Boxes delineate the regions used to assign dosage-inferred genotypes (DI-genotypes) based on their expected combinations of relative depth and allelic coverage fraction. For example, the standard diploid heterozygous genotype AB is expected at 2X depth and an allelic coverage fraction of 0.5, whereas AAB and AAAB are expected at 3X and 1/3 and at 4X and 1/4, respectively. The oval indicates a region of increased SNP density that does not correspond to an expected DI-genotype of duplication-associated SNPs and may result from mapping bias or somatic aneuploidy.

**Supplementary Figure 10.**
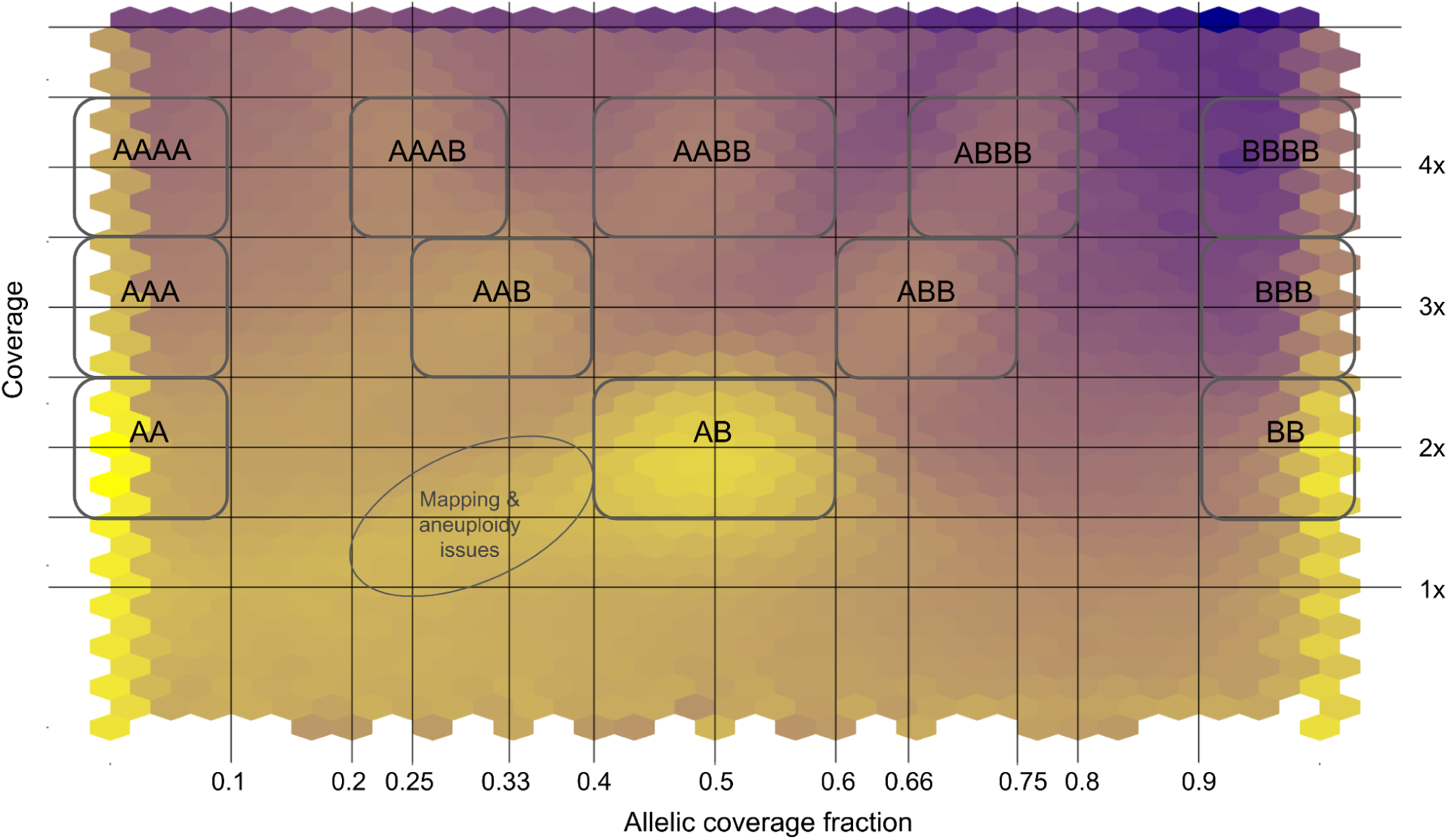
Density plot for sample 6C9M showing the joint distribution of allelic coverage fraction (x-axis) and relative sequencing depth (y-axis). Relative depth corresponds to SNP coverage normalized by the estimated haploid genome coverage, with the expected CN2, CN3 and CN4 depths indicated as 2X, 3X and 4X, respectively. Higher SNP densities are shown in brighter yellow. Boxes delineate the regions used to assign dosage-inferred genotypes (DI-genotypes) based on their expected combinations of relative depth and allelic coverage fraction. For example, the standard diploid heterozygous genotype AB is expected at 2X depth and an allelic coverage fraction of 0.5, whereas AAB and AAAB are expected at 3X and 1/3 and at 4X and 1/4, respectively. The oval indicates a region of increased SNP density that does not correspond to an expected DI-genotype of duplication-associated SNPs and may result from mapping bias or somatic aneuploidy.

**Supplementary Figure 11.**
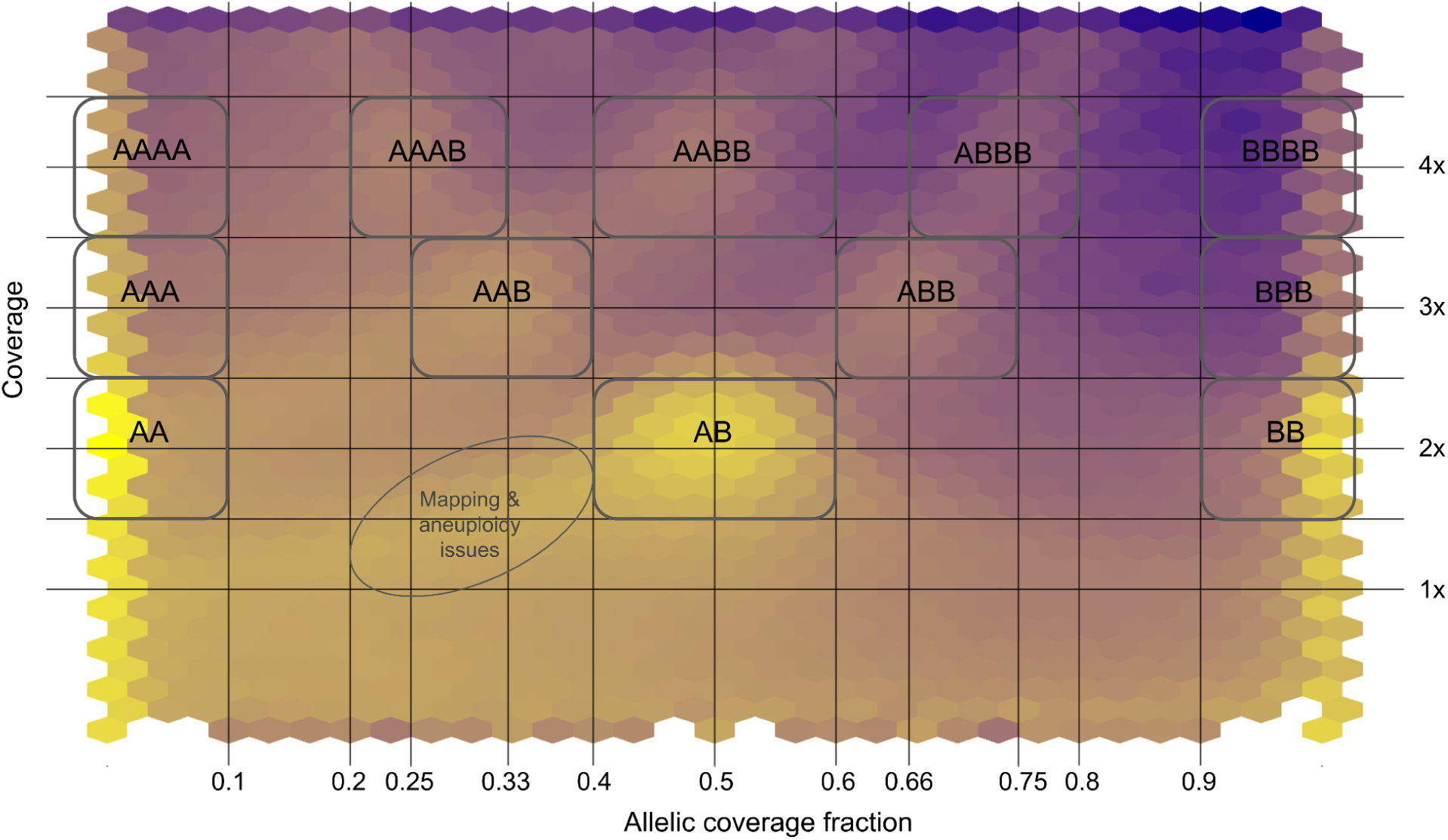
Density plot for sample 8C16M showing the joint distribution of allelic coverage fraction (x-axis) and relative sequencing depth (y-axis). Relative depth corresponds to SNP coverage normalized by the estimated haploid genome coverage, with the expected CN2, CN3 and CN4 depths indicated as 2X, 3X and 4X, respectively. Higher SNP densities are shown in brighter yellow. Boxes delineate the regions used to assign dosage-inferred genotypes (DI-genotypes) based on their expected combinations of relative depth and allelic coverage fraction. For example, the standard diploid heterozygous genotype AB is expected at 2X depth and an allelic coverage fraction of 0.5, whereas AAB and AAAB are expected at 3X and 1/3 and at 4X and 1/4, respectively. The oval indicates a region of increased SNP density that does not correspond to an expected DI-genotype of duplication-associated SNPs and may result from mapping bias or somatic aneuploidy.

**Supplementary Figure 12.**
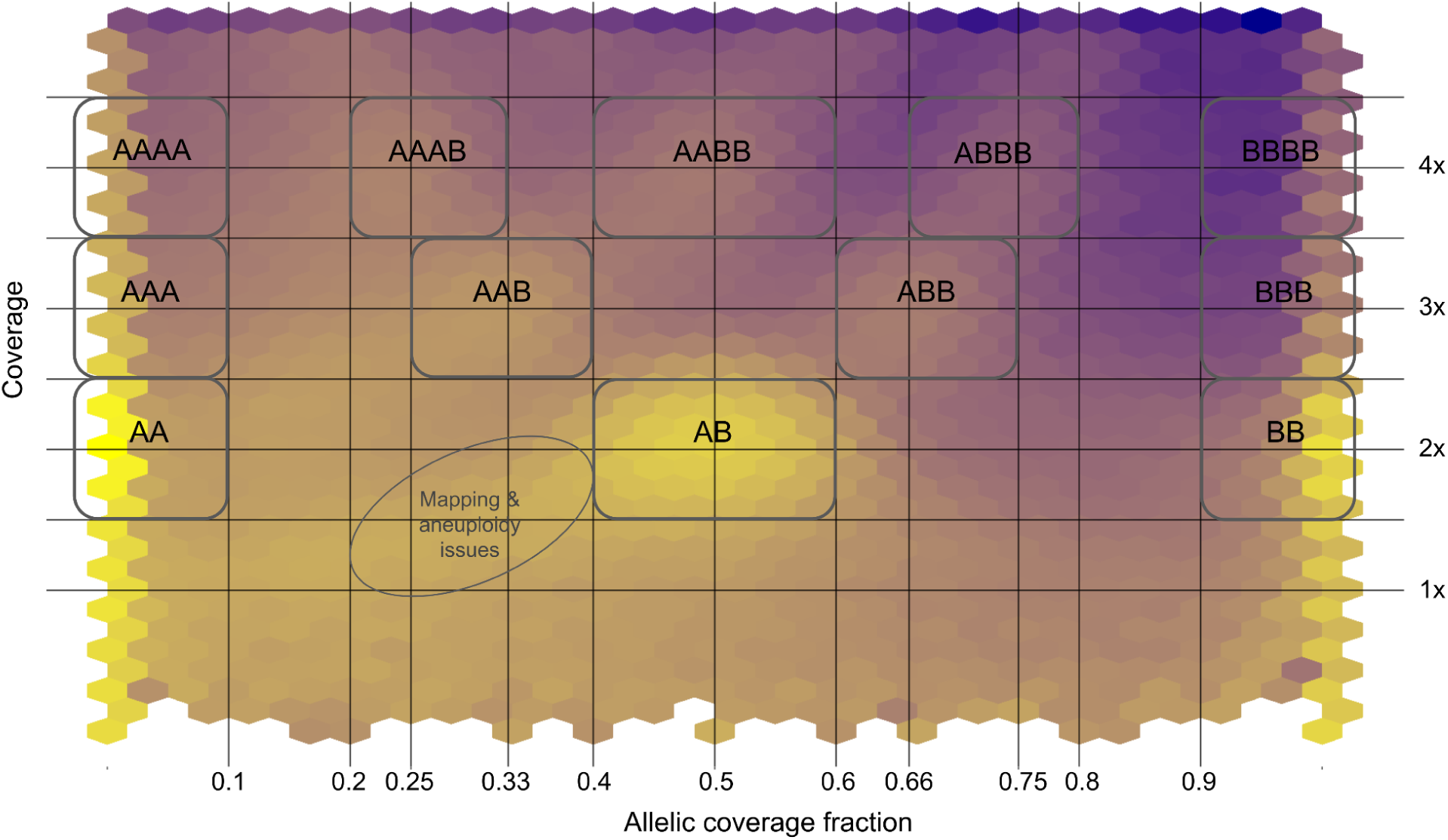
Density plot for sample ROSC showing the joint distribution of allelic coverage fraction (x-axis) and relative sequencing depth (y-axis). Relative depth corresponds to SNP coverage normalized by the estimated haploid genome coverage, with the expected CN2, CN3 and CN4 depths indicated as 2X, 3X and 4X, respectively. Higher SNP densities are shown in brighter yellow. Boxes delineate the regions used to assign dosage-inferred genotypes (DI-genotypes) based on their expected combinations of relative depth and allelic coverage fraction. For example, the standard diploid heterozygous genotype AB is expected at 2X depth and an allelic coverage fraction of 0.5, whereas AAB and AAAB are expected at 3X and 1/3 and at 4X and 1/4, respectively. The oval indicates a region of increased SNP density that does not correspond to an expected DI-genotype of duplication-associated SNPs and may result from mapping bias or somatic aneuploidy.

**Supplementary Figure 13.**
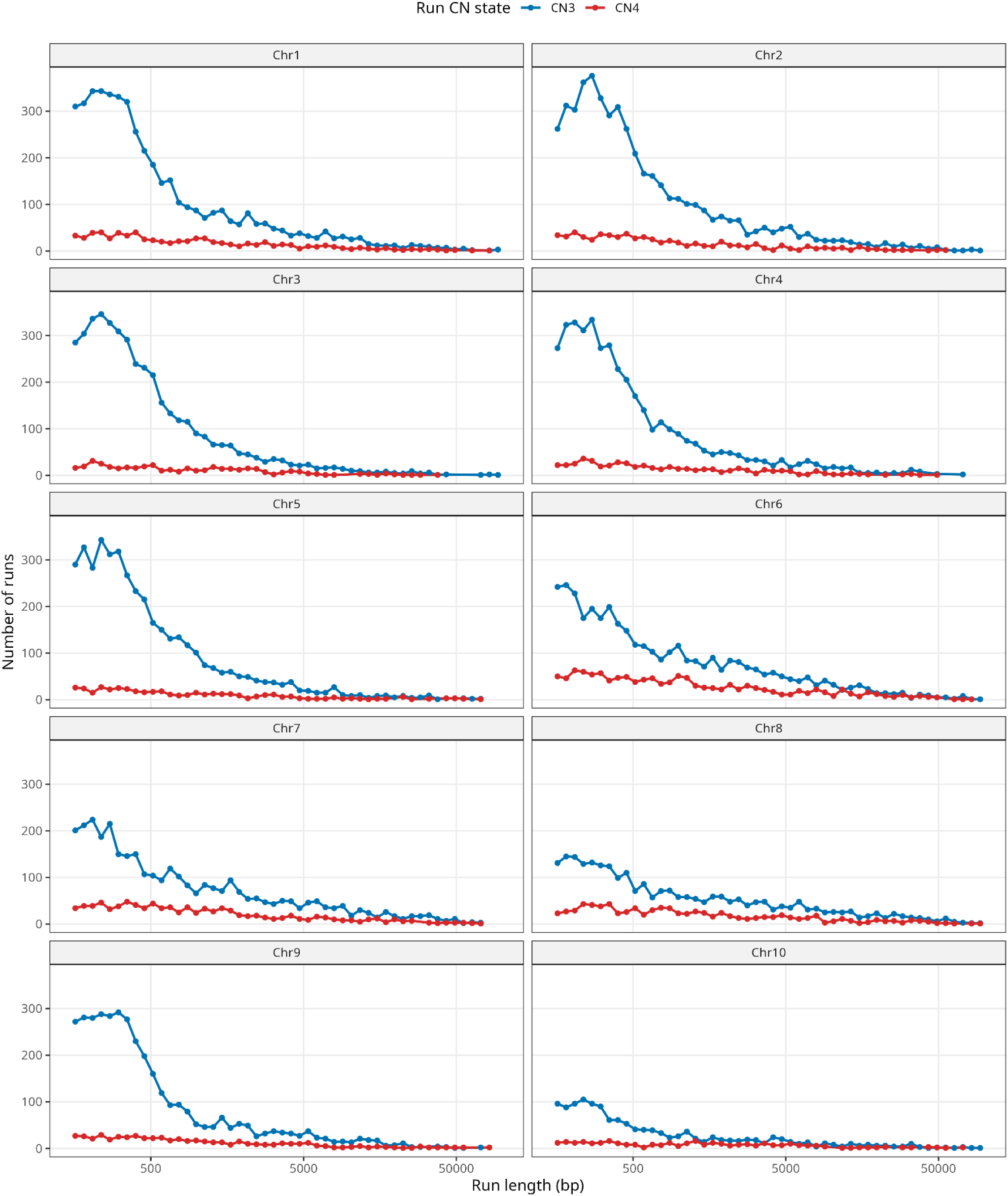
Distribution of CN3 and CN4 run lengths across chromosomes. Runs were defined as consecutive SNPs assigned to the same CN3 or CN4 dosage category. CN3 runs were more frequent than CN4 runs, but both showed similar distributions.

**Supplementary Figure 14.**
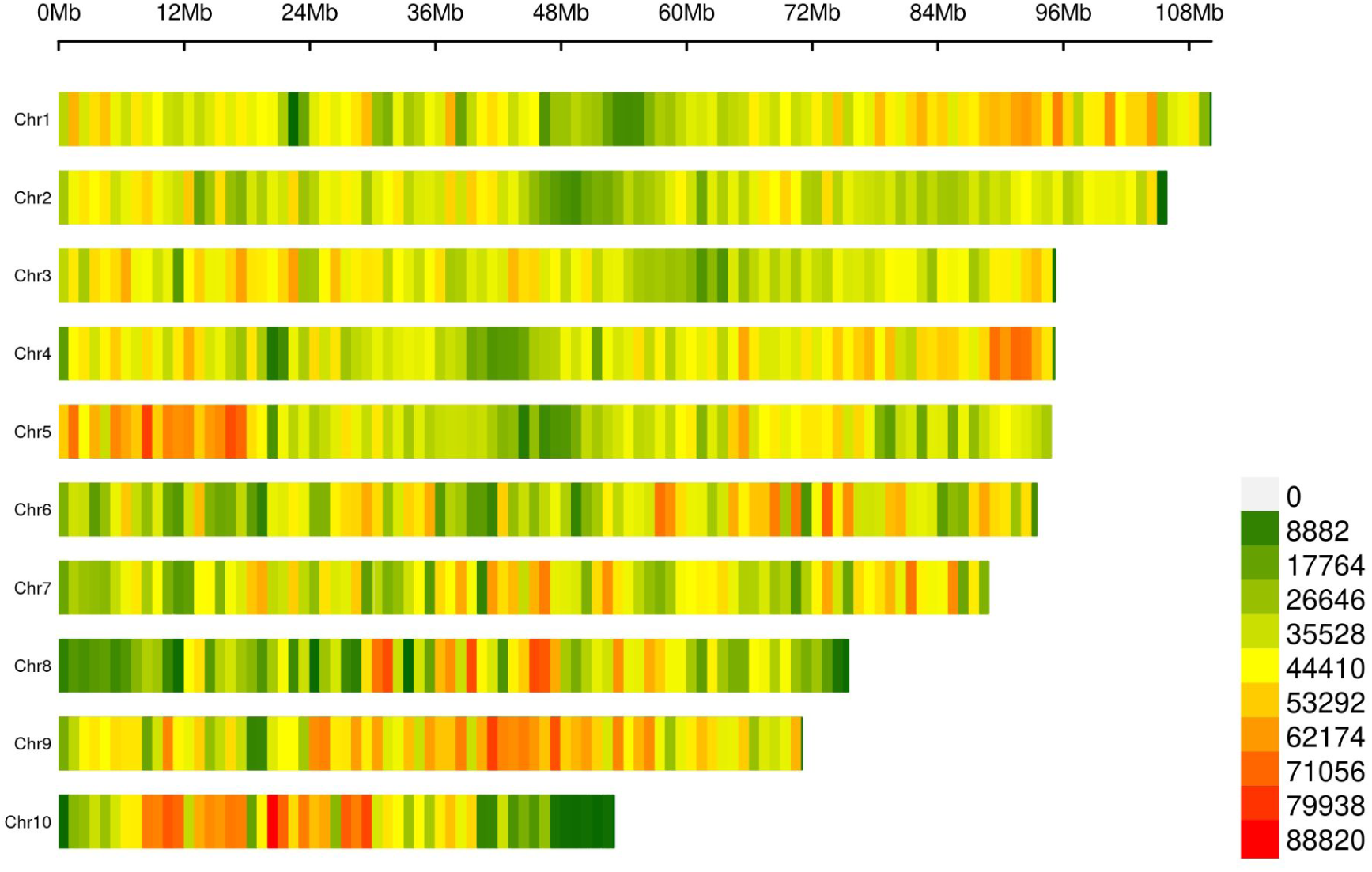
SNP-density plots averaged across all five individuals for 2X SNPs. Density of SNPs per 1Mb window shown for all 10 chromosomes. Position on the chromosome indicated on the x-axis. Note that regions of increased density at the end of chromosome 4, the start of chromosome 5, and second half of chromosome 9 correspond to known polymorphic inversions in this species (Lapègue et al. 2023, Sambade et al. 2022, Alves Monteiro et al. 2024, and see Supplementary Table 3).

**Supplementary Figure 15.**
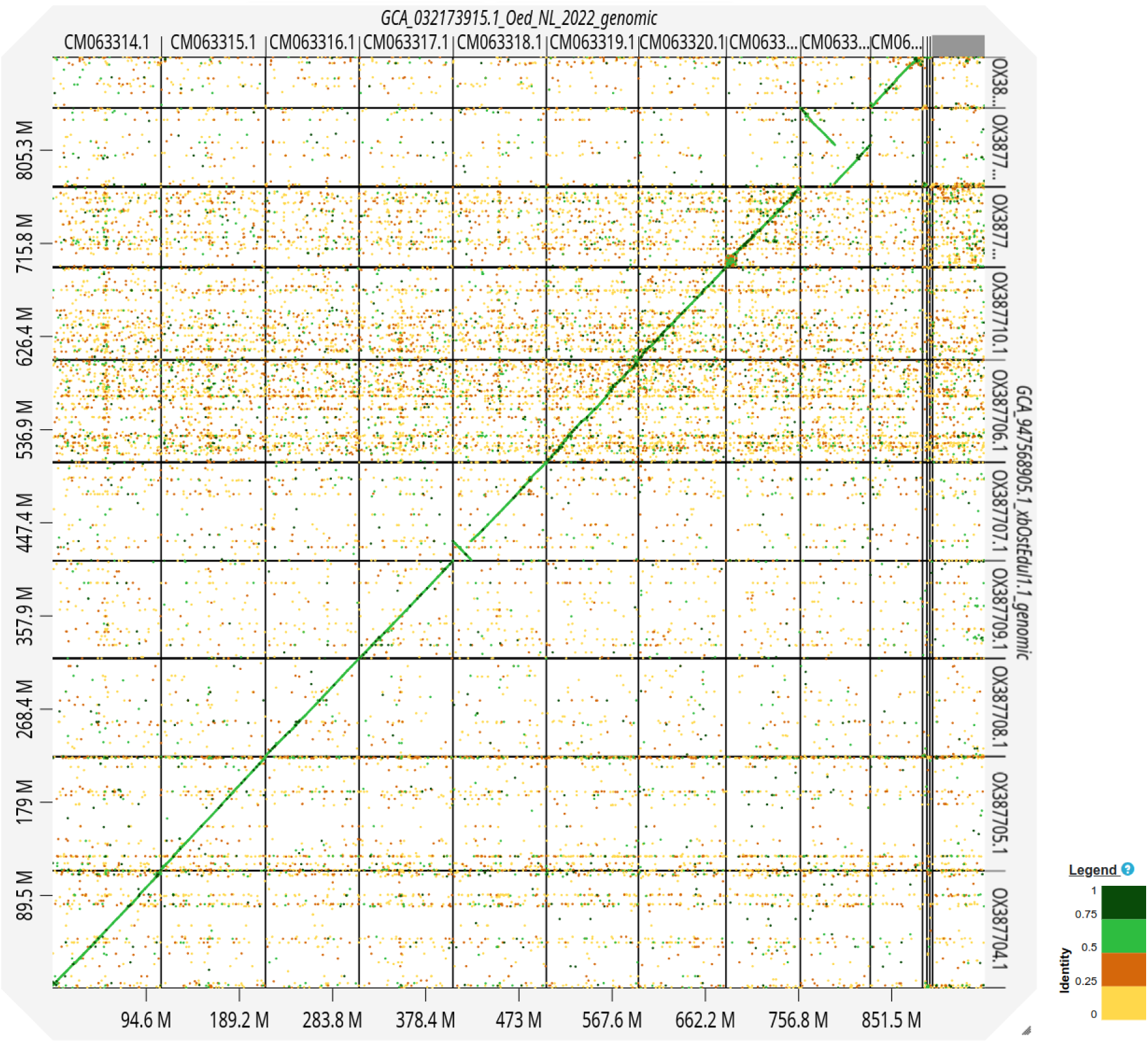
Whole-genome alignment between two Ostrea edulis reference assemblies visualized with D-GENIES. The Oed_NL_2022 assembly (Li et al. 2023), used as the reference genome in this study, is shown on the x-axis, and the Sanger Institute xbOstEdul1.1 assembly on the y-axis. Alignment patterns illustrate chromosome-scale synteny and differences in repeat content among chromosomes. The same comparison can also be viewed using NCBI’s Comparative Genome Viewer by following this URL: https://www.ncbi.nlm.nih.gov/cgv/browse/GCA_032173915.1/GCF_947568905.1/86715/37623

## SUPPLEMENTARY STATISTICS

### χ 2 test for uncharacterized/characterized SNPs in chromosomes 6, 7 and 8

Pearson’s Chi-squared test completed in base R using chisq.test

### 2C4M

X-squared = 183120, df = 2, p-value < 2.2e-16

### 4C6M

X-squared = 28858, df = 2, p-value < 2.2e-16

### 6C9M

X-squared = 181402, df = 2, p-value < 2.2e-16

### 8C17M

X-squared = 159831, df = 2, p-value < 2.2e-16

### ROSC

X-squared = 145161, df = 2, p-value < 2.2e-16

### χ 2 test for functional categories of SNPs classified as 2X, 3X and 4X SNPs

Pearson’s Chi-squared test completed in base R using chisq.test

### 2C4M

X-squared = 12147, df = 2, p-value < 2.2e-16

### 4C6M

X-squared = 22025, df = 2, p-value < 2.2e-16

### 6C9M

X-squared = 10706, df = 2, p-value < 2.2e-16

### 8C17M

X-squared = 14201, df = 2, p-value < 2.2e-16

### ROSC

X-squared = 10491, df = 2, p-value < 2.2e-16

